## Supplemental Figures and Tables for "*Dicer-like 5* deficiency confers temperature-sensitive male sterility in maize"

**Inventory of Supplementary Materials:**

Supplementary Figures 1-17

Supplementary Tables 1-3

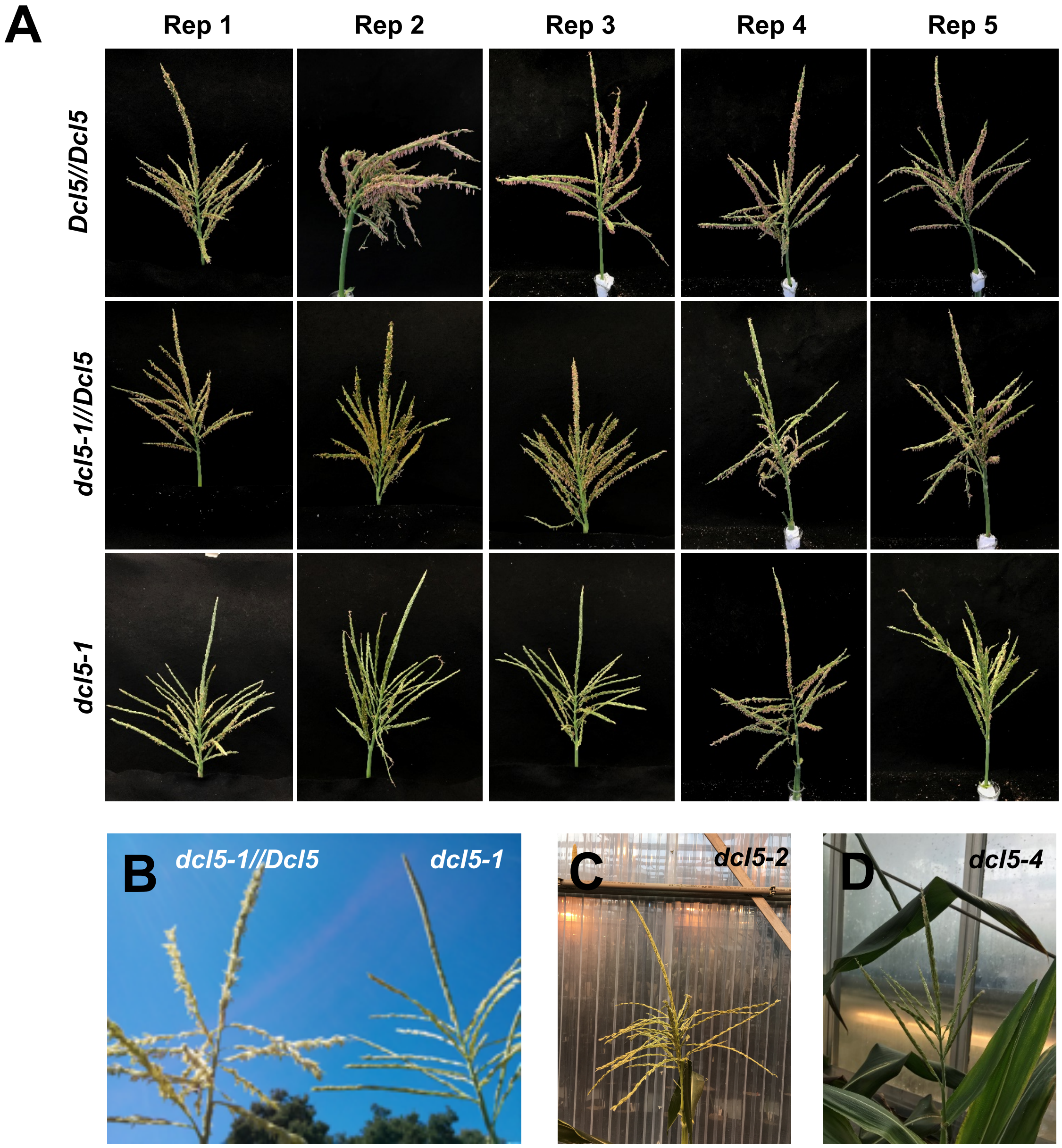


**Supplementary Figure 1. *dcl5-1* co-segregates with the male sterility phenotype.**

**(A)** In the *dcl5-1* segregating family, the male sterility phenotype co-segregates with the *dcl5-1* allele. In example photos shown above, all the wildtype *Dcl5//Dcl5* and heterozygous *dcl5-1//Dcl5* individuals were fertile but *dcl5-1* homozygous individuals were predominantly male sterile. **(B)** Male sterility phenotype of *dcl5-1* (right) in the field compared to its fertile sibling (left). **(C)** Male sterility phenotype of *dcl5-2*. **(D)** Male sterility phenotype of *dcl5-4*.

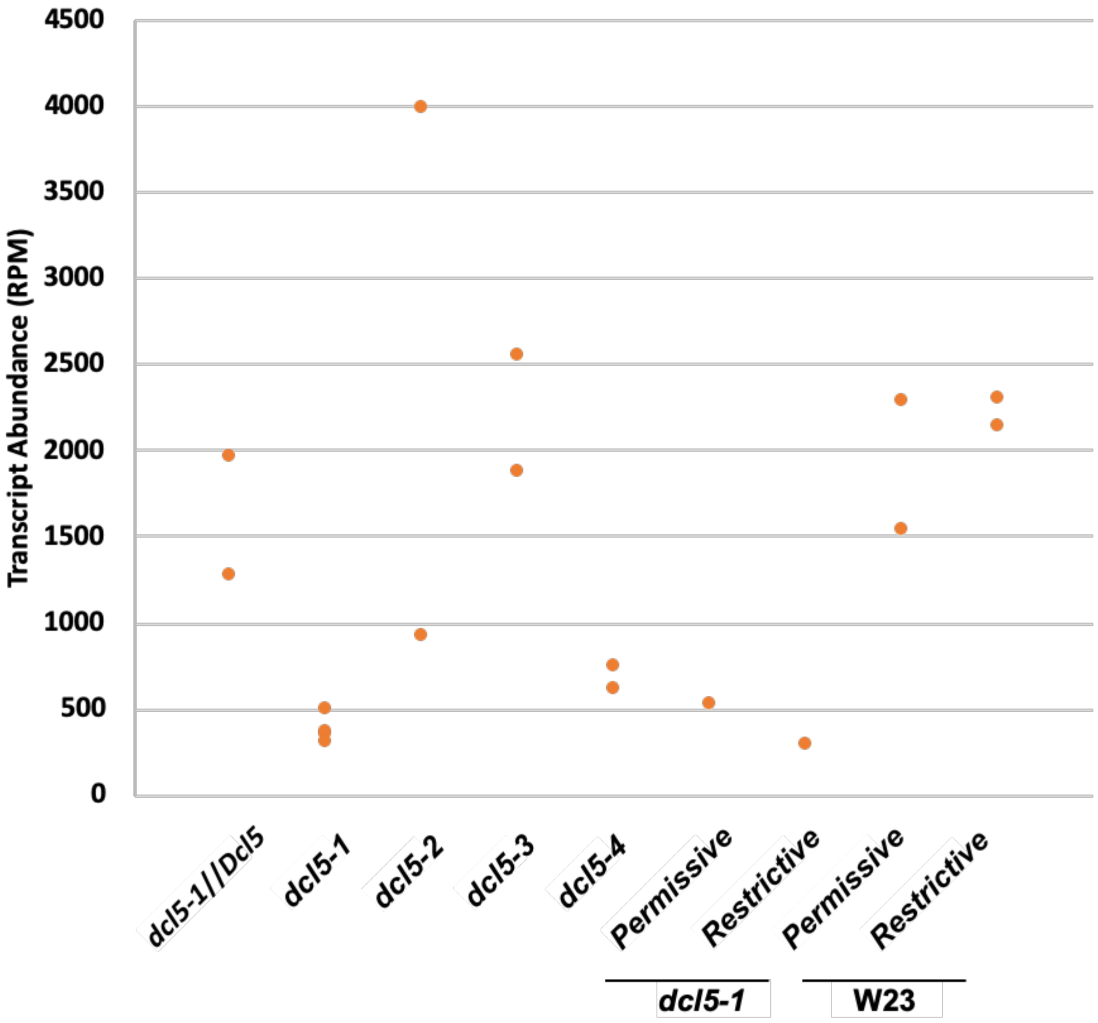


**Supplementary Figure 2. *Dcl5* mRNA levels in 2.0 mm anthers of control and mutant plants.**

*Dcl5* transcript levels retrieved from RNA-seq data are indicated for each genotype. *dcl5-1* and *dcl5-4* are frameshift mutants with transcripts levels about a third of their wild type siblings at 2.0 mm. In contrast, *dcl5-2* and *dcl5-3* contain short in-frame deletions and have similar or higher transcript levels compared to control (*dcl5-1*//*Dcl5*). For the *dcl5-1* plants grown under "restrictive" conditions, which exhibit developmental differences, spikelets equivalent to 2.0 mm anthers were analyzed. They contain the same transcript levels as the other *dcl5-1* samples under the field and "permissive" conditions. *Dcl5* transcripts levels in 2.0 mm anthers of the W23 inbred line are not significantly different under "restrictive" and "permissive" conditions.

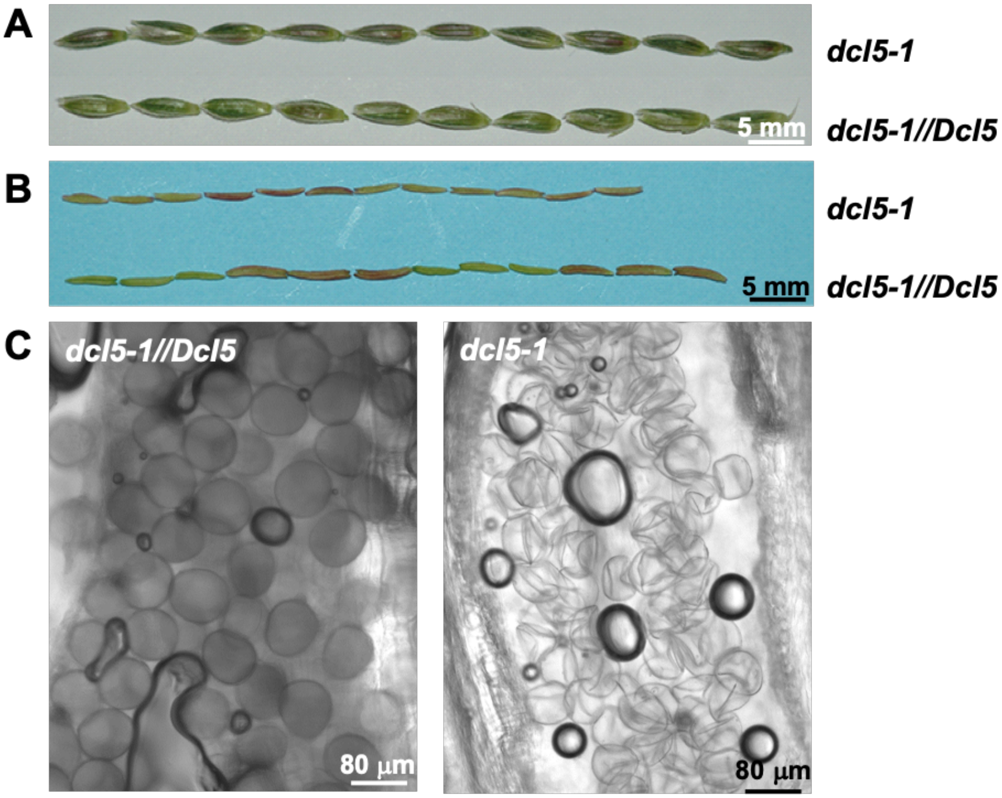


**Supplementary Figure 3.** ***dcl5-1*** **plants have normal spikelets but shorter, sterile anthers.**

*dcl5-1* samples are from homozygous, sterile plants; *dcl5-1//Dcl5* is a fertile sibling. **(A)** Dissected nearly-mature spikelets are equivalent in fertile and sterile samples. **(B)** Sterile anthers selected from the upper floret of spikelets in **(A)** are about 10% shorter than fertile anthers. **(C)** Pollen grains in *dcl5-1* anthers (right) are clear and collapsed relative to opaque and viable pollen from fertile anthers (left). Bars in **(A, B)**, 5 mm; bars in **(C, D)**, 80 μm.

**A**

```
Guide RNA 5′-GTTCATGGTCTATCTGGAAACTGG...76 nt...CTGGGTTCGTTCTCTTAATGG-3′
Translated    V  H  G  L  S  G  N                  G  F  V  L  L  M
```

**B**

```
ZmDCL1   EGVAEILHGEWILSGRDGYQSSQFIKLYLYSVNCVNIGTSKDPFVTQLSNFALIFGNELD
ZmDCL4   MLIPAVLRPSRCKLDSS-----FKLHFYYIEFIPIPADR-------RYQMFGLFVINRLP
ZmDCL2   DYFPEQLVDNWSSFSRR-----GFYYCY----MVSLEGCSK----TTPAEIVLAVKCDMG
ZmDCL3   TTKVLSMSGSWGSDRSV-----TKLQGYKLNFVCDQVGQK-------YSAFVLLIDASIA
ZmDCL5   TINVHGLSGNWIHESET-----VILNAYTFDFRCDQEGEL-------YAGFVLLMESVLD
               :  .                 *         .            : * .   :
```

**C**

```
ZmDCL3   ------TACAACCAAAGTTCTTTCCATGTCTGGCTCTTGGGGTTCAG--
ZmDCL5   ------AACGATAAATGTTCATGGTCTATCTGGAAACTGGATCCATG--
ZmDCL2   --------TGACTACTTTCCAGAGCAATTAGTGGATAATTGGTCGTCCT
ZmDCL4   GATGTTAATCCCTGCAGTTCTGAGACCTTCAAGATGCAAAC--------
                          * *        *   *
```

**D**

```
ZmDCL2   ------AACTACTCCAGCTGAGATAGT—ATTAGCTGTAAAGTGCGACA
ZmDCL3   --TCAAAAATACTCTGCCTTTGTCCTTTTAATTGATGCTAGTA-----
ZmDCL5   GGTGAACTTTATGCTGGGTTCGTTCTCTTAATGGAATCAGT-------
ZmDCL1   --TACACAACTTTCAAATTTTGCTCTT--ATTTTTGGCAATGAGC---
ZmDCL4   --TAGACGATATCAGATGTTTGGTCTT--TTTGTGATCAATCGCC---
                           *  *         *
```

**E**

```
ZmDCL3   TACAACCAAAGTTCTTTCCATGTCTGGCTCTTGGGGTTCAG
ZmDCL5   AACGATAAATGTTCATGGTCTATCTGGAAACTGGATCCATG
          ** *  ** **** *    * *****    ***

ZmDCL3   TCAAAAATACTCTGCCTTTGTCCTTTTAATTGATGCTAG
ZmDCL5   TGAACTTTATGCTGGGTTCGTTCTCTTAATGGAATCAGT
         * **   **  ***  ** ** ** ***** **  *
```

**Supplementary Figure 4. CRISPR-Cas9 gRNAs targeting maize *Dcl5* lack homology to *Dcl3***

**(A)** Guide RNA sequences of *Dcl5* (upper line, magenta letters), PAM sequences (upper line, green letters), and the corresponding translated sequences (lower line, magenta letters). **(B)** Translated sequence alignment of the maize *Dcl* genes, by CLUSTAL O (1.2.4) multiple sequence alignment. The region including the translated guide RNA sequences is shown here. The translated guide RNA sequences of *Dcl5* are highlighted in magenta and the non-conserved residues in the corresponding sequences of the other *Dcl* genes are in green. The first **(C)** and second **(D)** guide RNA sequences targeting *Dcl5* are highlighted in magenta, PAM sequences are highlighted in green, and the corresponding sequences of the other *Dcl* genes are aligned by CLUSTAL O (1.2.4) multiple sequence alignment. The corresponding sequences of the first guide RNA in *Dcl1* is split in two exons and is not included in **(C)**. **(E)** Although *Dcl3* and *Dcl5* are highly conserved, the corresponding sequence in *Dcl3* could not be targeted by either guide RNA designed for *Dcl5*. In the first gRNA (upper) there is poor sequence similarity, and the PAM sequence (green letters) of the second gRNA (lower) is not present in *Dcl3*. An * (asterisk) indicates positions which have a single, fully conserved residue; a colon indicates conservation between groups of strongly similar properties, scoring > 0.5 in the Gonnet PAM 250 matrix; a dot or period indicates conservation between groups of weakly similar properties, scoring ≤ 0.5 in the Gonnet PAM 250 matrix.

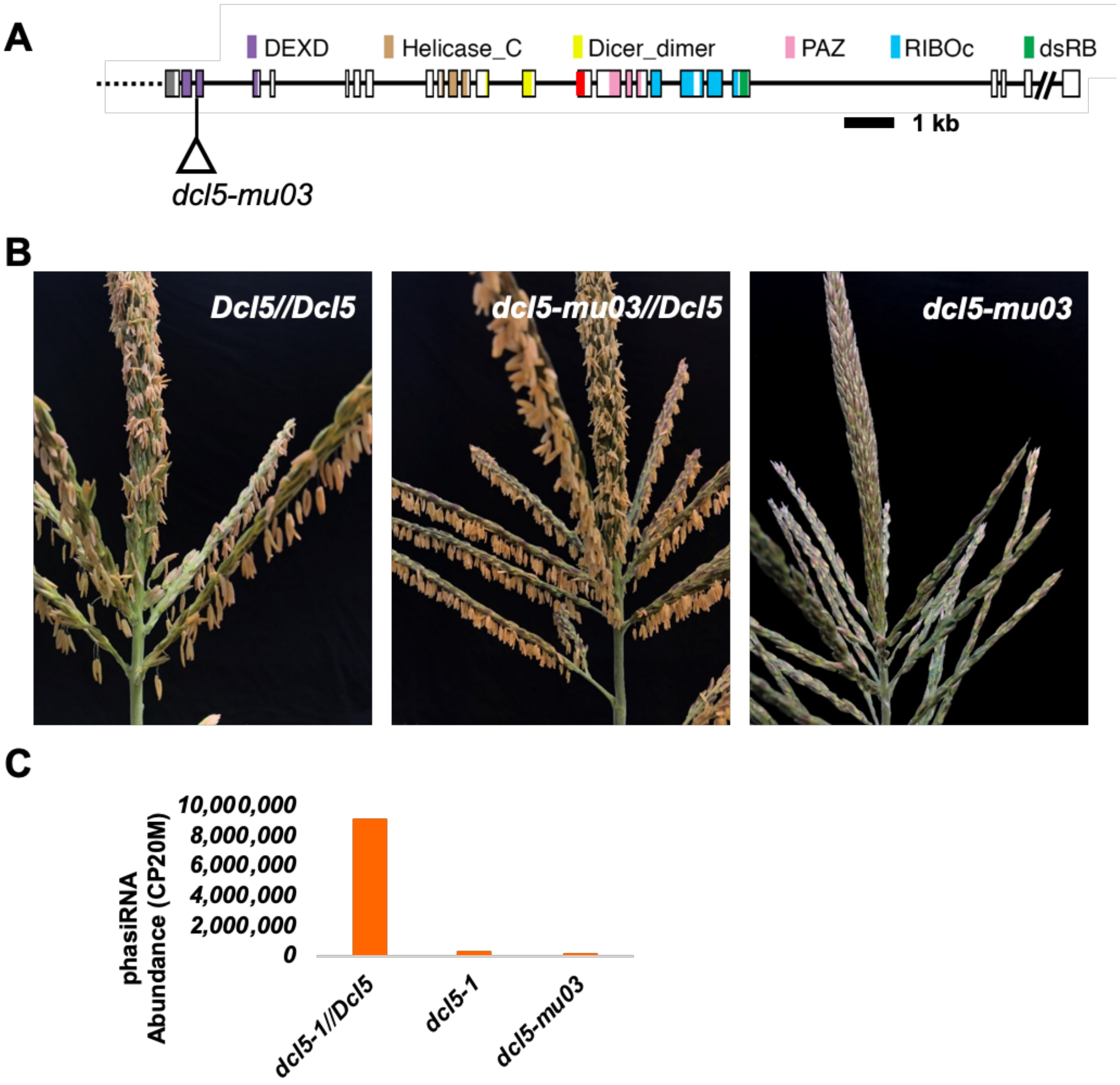


**Supplementary Figure 5. A *Mu* insertion allele of *Dcl5* lacks 24-nt phasiRNAs and is male sterile.**

**(A)** Schematic diagram of the *Dcl5* gene model. *dcl5-mu03* has a *Mu* transposon insertion in the third coding exon, yielding a null allele. **(B)** A *dcl5-mu03* tassel lacking any exerted anthers (right) and sibling *dcl5-mu03//Dcl5* and *Dcl5//Dcl5* plants at peak pollen shed with hundreds of exerted anthers (left and middle). **(C)** Near absence of 24-nt phasiRNAs in 2.0 mm anthers of *dcl5-mu03* mutants. This result is highly similar to the case for *dcl5-1* in comparison to the heterozygous siblings.

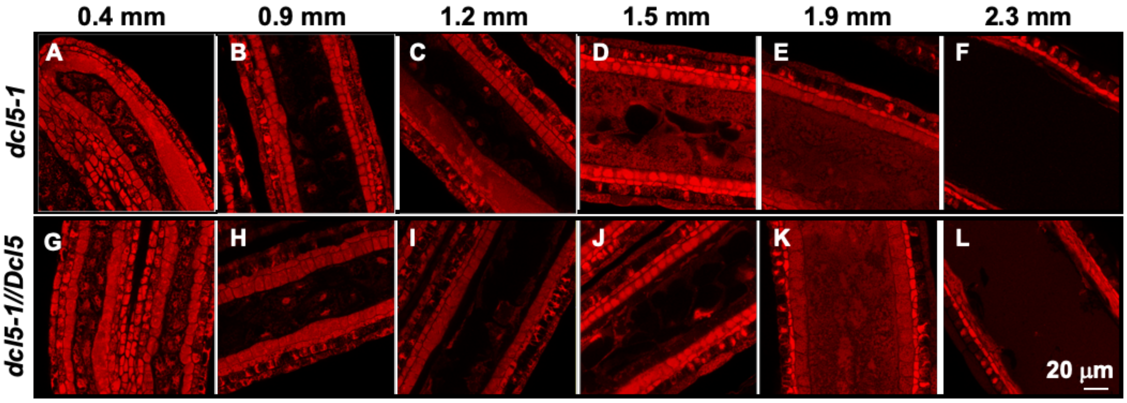


**Supplementary Figure 6. Periclinal divisions establish normal cell layers and cell morphology in *dcl5-1* and *dcl5-1//Dcl5* control anthers with the same developmental timing.**

Anthers of 0.4 mm, 0.9 mm, 1.2 mm, 1.5 mm, 1.9 mm, and 2.3 mm were dissected from homozygous mutant and heterozygous *dcl5-1//Dcl5* sibling plants, fixed, stained with propidium iodide, and observed by confocal microscopy. Twenty-three anthers from five *dcl5-1//Dcl5* and thirty-five anthers from eight *dcl5-1* plants, ranging from 0.3 mm to 2.3 mm were used in this study. No obvious defects in cell layer differentiation were observed in *dcl5-1* anthers **(A-F)** in comparison to their control siblings **(G-L)**. Plants were grown under a temperature regime that causes male sterility in *dcl5-1.* The scale bar in **(L)** is approximate and pertains to all the other panels, 20 μm.

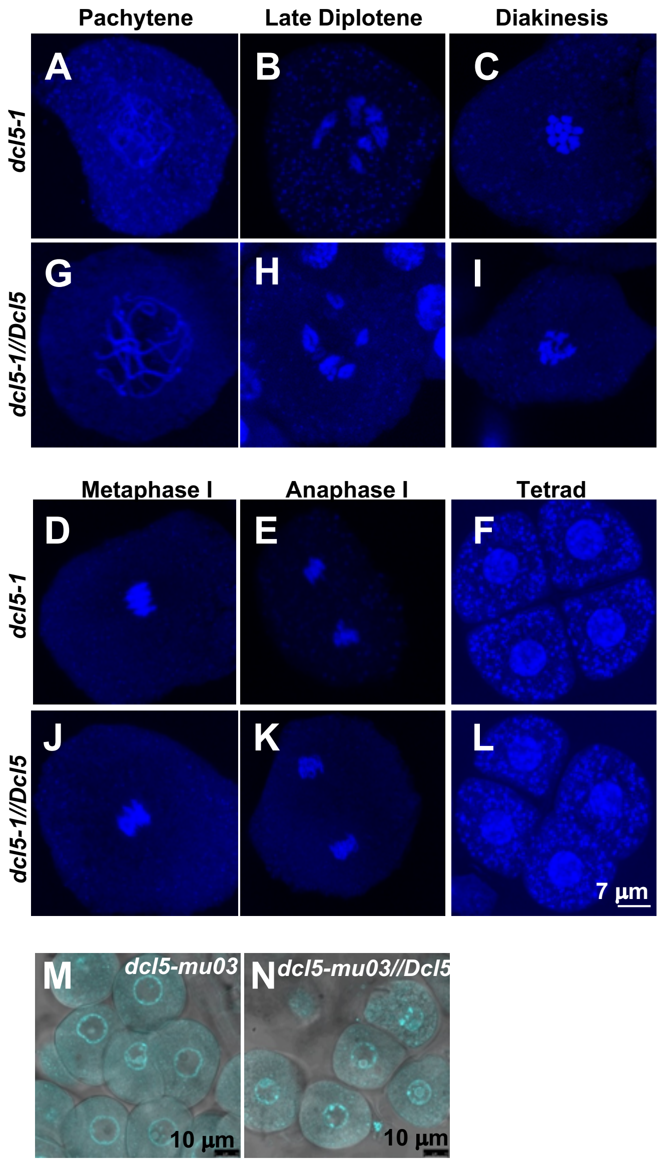


**Supplementary Figure 7. Meiotic progression in *dcl5-1* and *dcl5-mu03* anthers was nearly normal.**

Meiocytes were extruded from homozygous mutant and heterozygous *dcl5-1//Dcl5* sibling plants, fixed, and stained with DAPI. In the majority of the meiotic cells, no obvious defects were observed during the periods of homolog search and chromosome pairing, chromosome alignment, and division in *dcl5-1* anthers **(A-F)** compared to their fertile siblings **(G-L)**. Meiosis progression in five *dcl5-1//Dcl5* and six *dcl5-1* plants were used in this study. Unlike previously reported defective meiosis in the rice *mel1* mutant, *dcl5-1* meiotic progression is normal. *dcl5-mu03* also produces uninucleate microspores **(M)**, with the same properties as in *dcl5-mu03//Dcl5* siblings **(N)**. Plants were grown under a temperature regime that causes male sterility in *dcl5* mutants. The scale bar in **(L)** is approximate and pertains to **(A-L)**, 7 µm; bars in **(M, N)**, 10 µm.

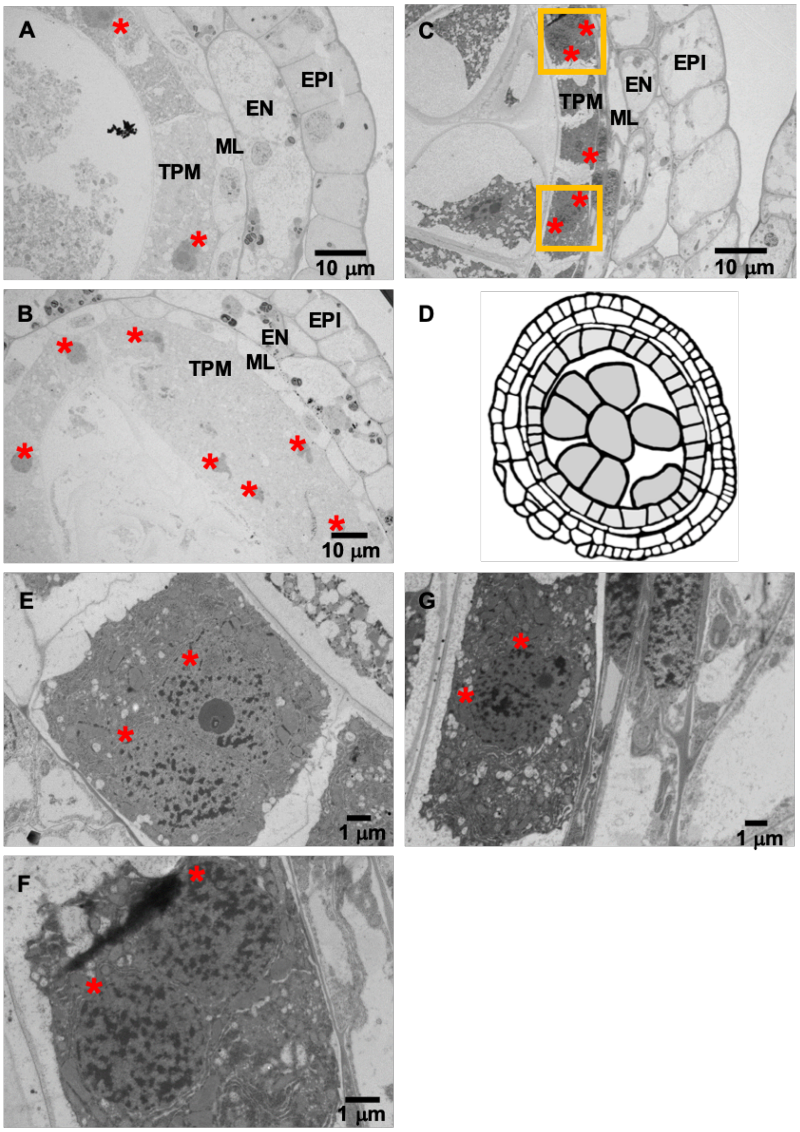


**Supplementary Figure 8. Tapetal cells are morphologically defective in *dcl5-1* anthers.**

Multiple transverse sections of *dcl5-1* 2.0 mm anthers (left) and *dcl5-1//Dcl5* anther (right) were prepared for TEM as indicated in the methods section. Both the mutant **(A-B)** and the fertile sibling **(C)** anther lobes contained five cell types: external epidermis, endothecium, middle layer, tapetum, and central meiocytes. Mutant tapetal cells are pale compared to the fertile sibling. A schematic of a ~2 mm anther is provided for reference **(D)**. Tapetal cell nuclei are indicated with red asterisks, highlighting the difference in proportion of binucleated tapetal cells between sterile mutant and fertile sibling anthers. **(E-G)** Enlarged binucleated tapetal cells in the boxes from Fig. 2A and Fig. S8C. Bars in **(A, B, C)**, 10 µm; Bars in **(E, F, G)**, 1 µm.

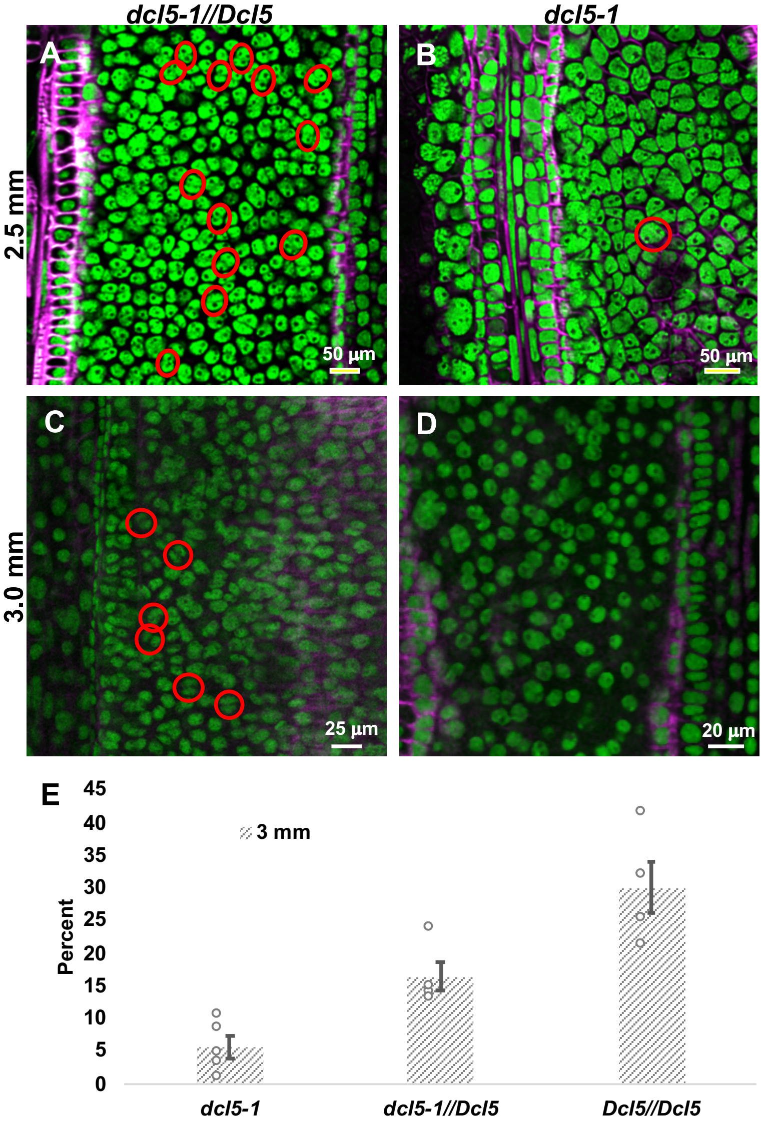


**Supplementary Figure 9. Substantially fewer binucleated tapetal cells in *dcl5-1* anthers compared to fertile anthers.**

Confocal imaging of the tapetal layer in cleared anthers using a range of anther sizes (2 to 3 mm) from *dcl5-1//Dcl5* **(A, C)** and *dcl5-1* **(B, D)** plants doubly stained with nuclear marker Syto13 (green) and cell wall marker Calcofluor white (pink). Example binucleated cells are circled in red.

**(A, B)** Binucleate tapetal cells at 2.5 mm.

**(C, D)** Binucleate tapetal cells at 3.0 mm.

**(E)** Quantification of binucleated tapetal cells in 3.0 mm anthers from *dcl5-1*, *dcl5-1//Dcl5*, and *Dcl5//Dcl5* siblings, in a family segregating 1:2:1. The binucleated cells were manually counted in 2-D images, leading to under counting of this class, because only one plane of the cell is in view. (p-value: *dcl5-1 comparing to dcl5-1//Dcl5, p = 0.00755; dcl5-1 comparing to Dcl5//Dcl5, p = 0.0008.*).

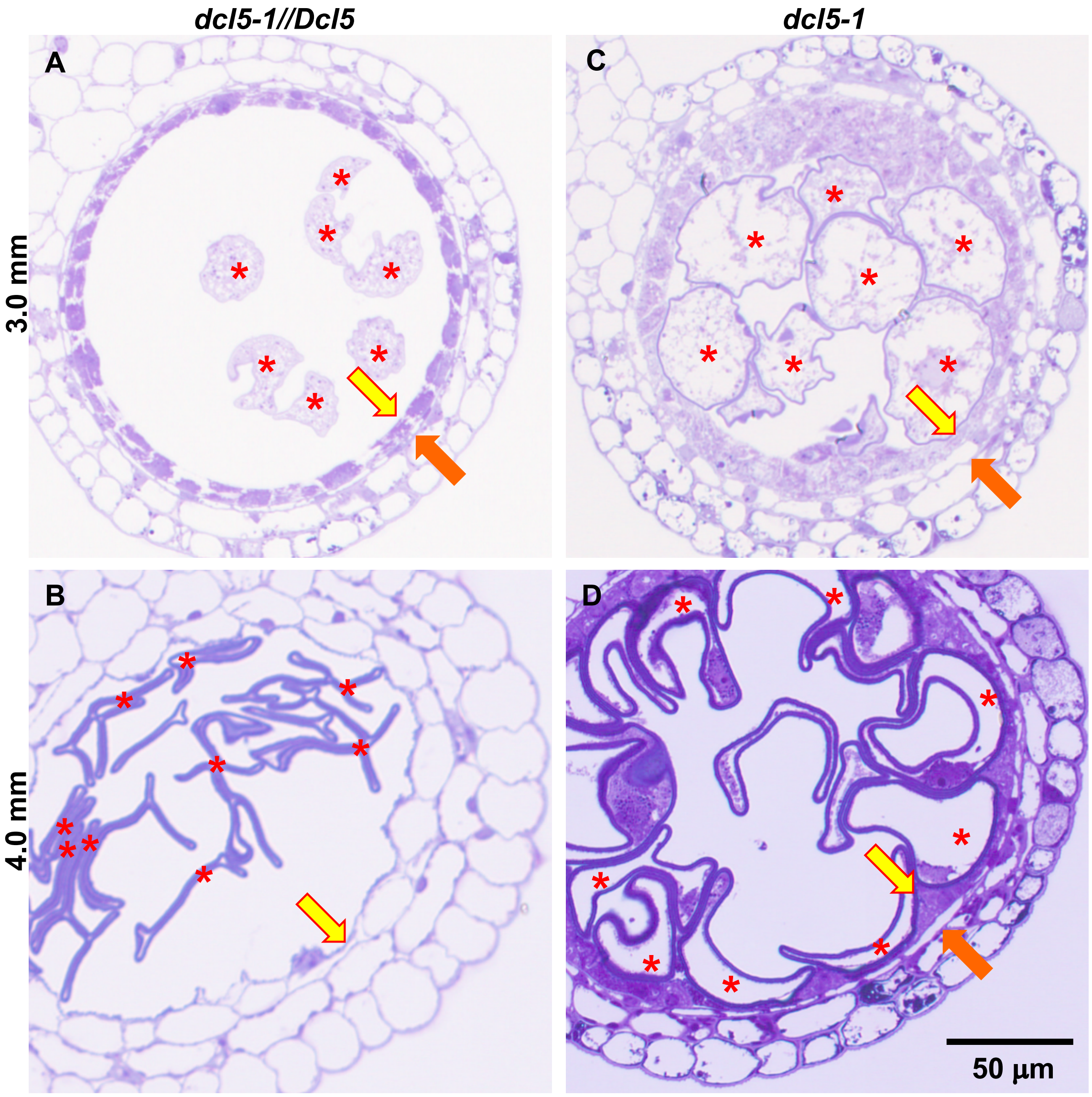


**Supplementary Figure 10. A delayed and retained tapetum in *dcl5-1*.**

Three replicates for each sample were studied. The tapetum initiates programmed cell death, becomes vacuolated, and ultimately collapses and disappears in 3.0 mm **(A)** and 4.0 mm **(B)** anthers of *dcl5-1//Dcl5* plants. In contrast, the tapetal layer is retained in the *dcl5-1* anthers at the 3.0 mm **(C)** and 4.0 mm **(D)** stages. Similarly, the middle layer is collapsed **(A)** and disappears **(B)** in these stages in *dcl5-1//Dcl5* anthers, but like the tapetum, it is retained in *dcl5-1* anthers **(C, D)**. Asterisks indicate unicellular microspores; yellow arrows, the tapetum or its remnants; orange arrows, middle layer. Bar in **(D)** is 50 µm and applies to all four images.

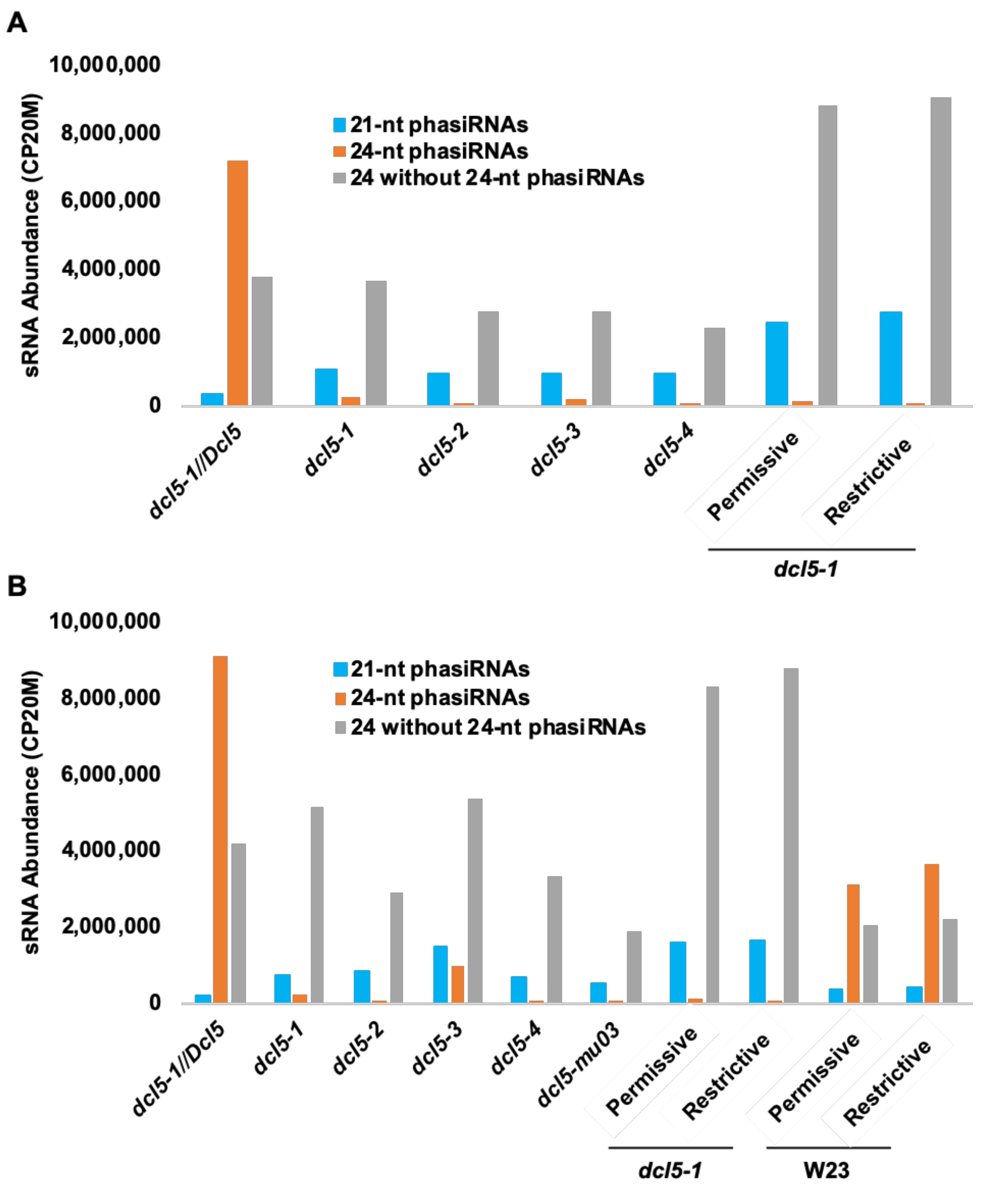


**Supplementary Figure 11. Among sRNA types, only abundances of 24-nt phasiRNAs were reduced in *dcl5* anthers.**

sRNA abundances (normalized to total reads) from 1.5 mm anthers **(A)** and 2.0 mm anthers **(B)**. Permissive refers to anthers dissected from plants grown at 21° C during meiosis and the subsequent six days; restrictive refers to anthers from plants grown at 28° C during this interval. The plants and libraries from *dcl5-1* and W23 under permissive and restrictive conditions were a separate experiment, and the increased levels of non-phased 24-nt sRNAs likely reflect different growth conditions and experimental variation.

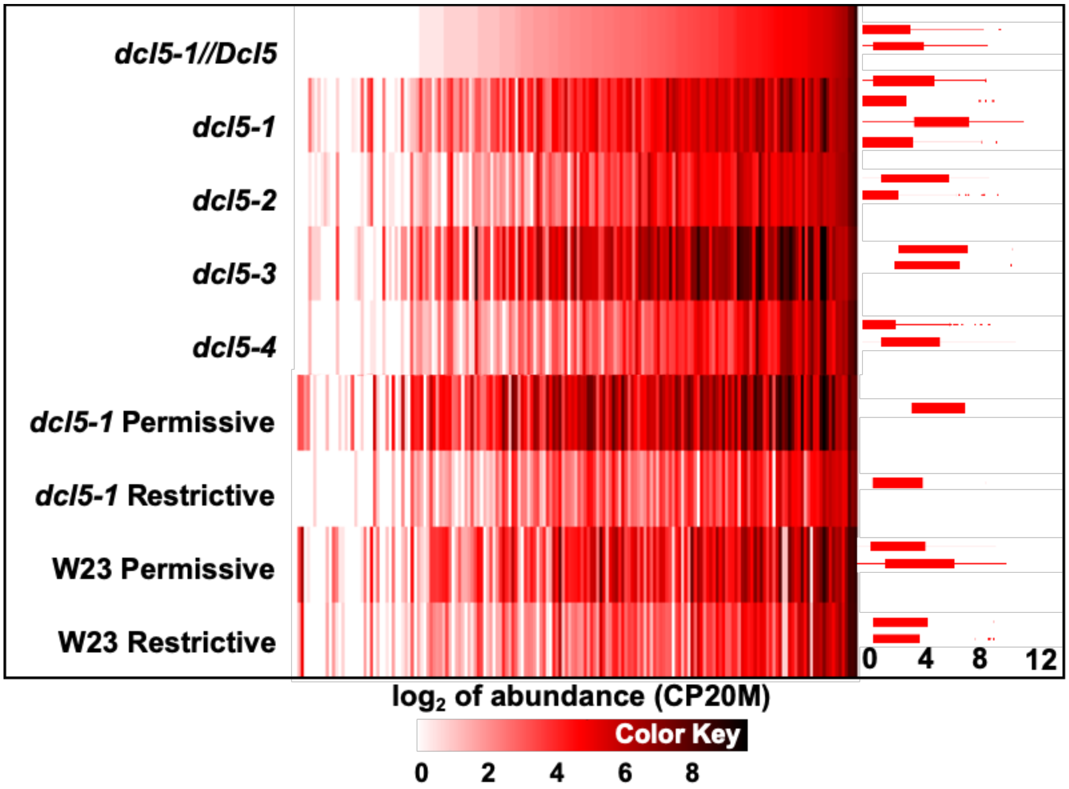


**Supplementary Figure 12.** ***24-PHAS*** **precursor accumulation in** ***dcl5*** **mutants.**
The RNA-seq reads from 2.0 mm anthers were mapped to 176 24-nt *PHAS* loci (vertical bars within each row); this analysis was performed for the heterozygous control (top row) and four homozygous *dcl5* mutants, as labeled. Replicates were averaged to generate a single abundance value for each allele. Shades of white to red to black represent the sum of hits-normalized abundances of RNA-seq reads, converted to $\log_2$ of abundance (see the key at the bottom). To the right, boxplots with the same scale are shown for the same group of *PHAS* loci, for each replicate allele; the center line represents the median, box limits are the upper and lower quartiles, whiskers are the 1.5x interquartile range, and points show the scatter of outliers.

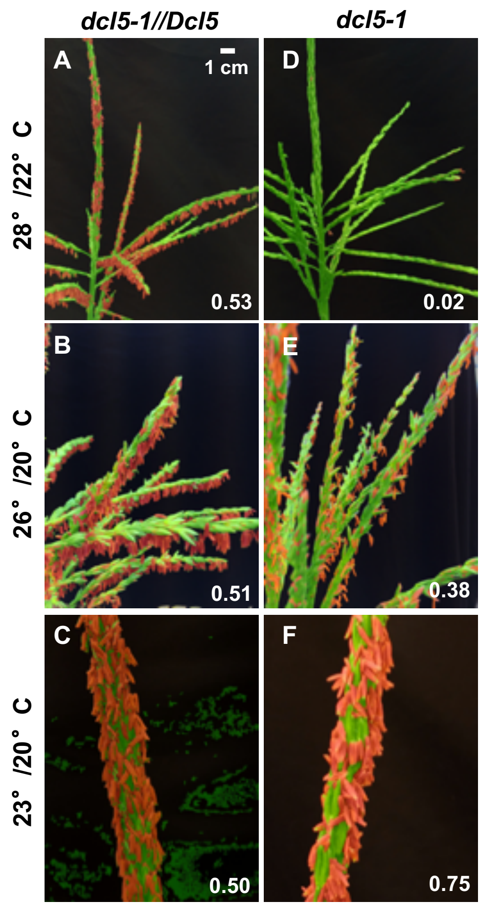


**Supplementary Figure 13.** ***dcl5-1*** **anther/tassel area ratio indicates** ***dcl5-1*** **is temperature sensitive.**

To better estimate the fertility of tassels under different temperature regimes, a naïve Bayes classifier-based machine learning method was applied to the images in Fig. 4. Pixels in each image were classified and labeled as anthers (red), the glum and branches of the tassel (green), and dark background (unmasked); then, a ratio representing pixels of anthers (red) to pixels of tassel (red and green, i.e. anthers + glum and branches) was determined for each image (number in white, bottom right). **(A, D)** Heterozygous *dcl5-1//Dcl5* and *dcl5-1* in a 28°/22°C regime; **(B, E)** Heterozygous *dcl5-1//Dcl5* and *dcl5-1* in 26°/20°C; **(C, F)** Heterozygous *dcl5-1//Dcl5* and *dcl5-1* in 23°/20°C. **(A, B, C)** Heterozygous *dcl5-1//Dcl5* siblings were fully fertile under all three regimes; the anther/tassel area ratio was about 0.50; **(D, E, F)** *dcl5-1* plants in the restrictive regime **(D)** were completely male sterile, and the ratio was 0.02; those in permissive conditions were partially **(E)** or fully **(F)** fertile, and the ratios were 0.38 and 0.75. The pace of tassel development is temperature-dependent and full anther exertion occurs at different days after planting, for example **(C)** was photographed on the first day of anther exertion, a seven-day process. In this case, we measured and compared main branches in **(C, F)**. The scale bar in **(A)** is approximate and pertains to all tassel images.

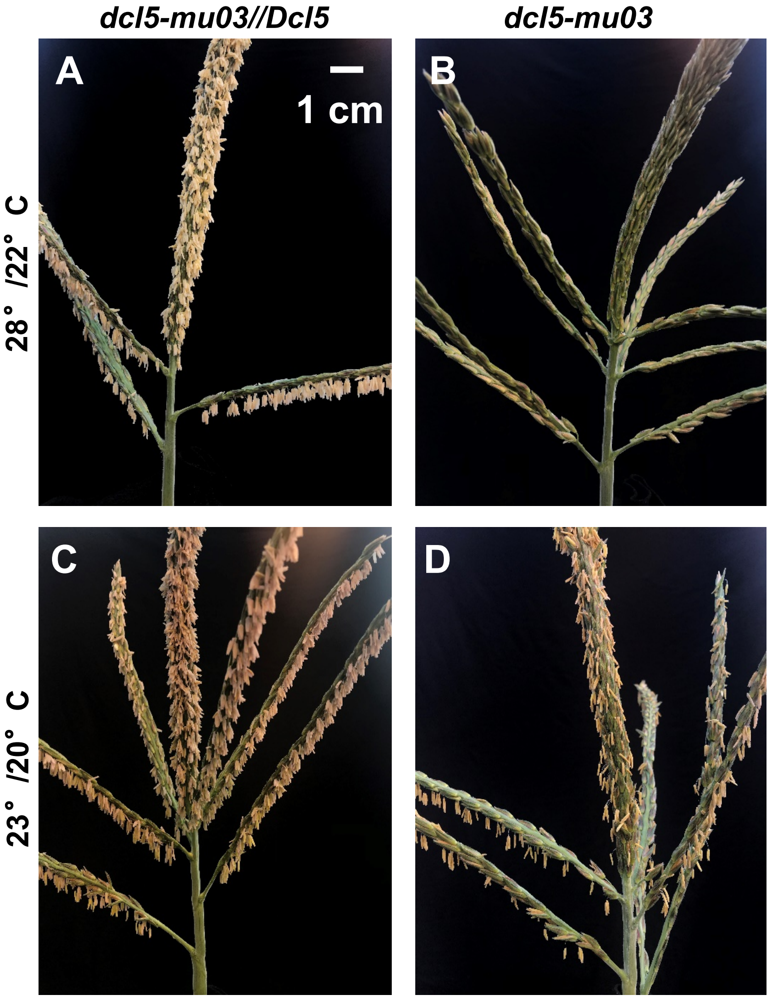


**Supplementary Figure 14.** ***dcl5-mu03*** **anther fertility is temperature sensitive.**

Two sets of *dcl5-mu03* and sibling *dcl5-mu03//Dcl5* plants grown in greenhouses with differing temperature regimes: 28°/22° C or 23°/20° C (day/night) from 40-57 day after planting (DAP); all the plants are from the same 3:1 segregating family and were grown at 28°/22° C for 1-40 DAP and after 57 DAP until pollen shed. **(A, C)** Heterozygous *dcl5-mu03//Dcl5* siblings were fully fertile under both regimes. **(B)** *dcl5-mu03* plants in the restrictive regime were completely male sterile, while those in permissive conditions were partially fertile **(D)**. There are at least 3 individual plants for each genotype in each regime as biological replicates, of which the recovered male fertility percentages correlated with flowering time – delayed flowering under cool permissive conditions correlated with higher anther exertion. Images were taken at 70±7 DAP; the pace of tassel development is temperature-dependent and full anther exertion occurs at different times, for example **(A, B)** was photographed ten days later after the first anther exertion, while **(C, D)** were photographed on the 3-5 days after the first anther exertion. The scale bar in **(A)** is approximate and pertains to all tassel images.

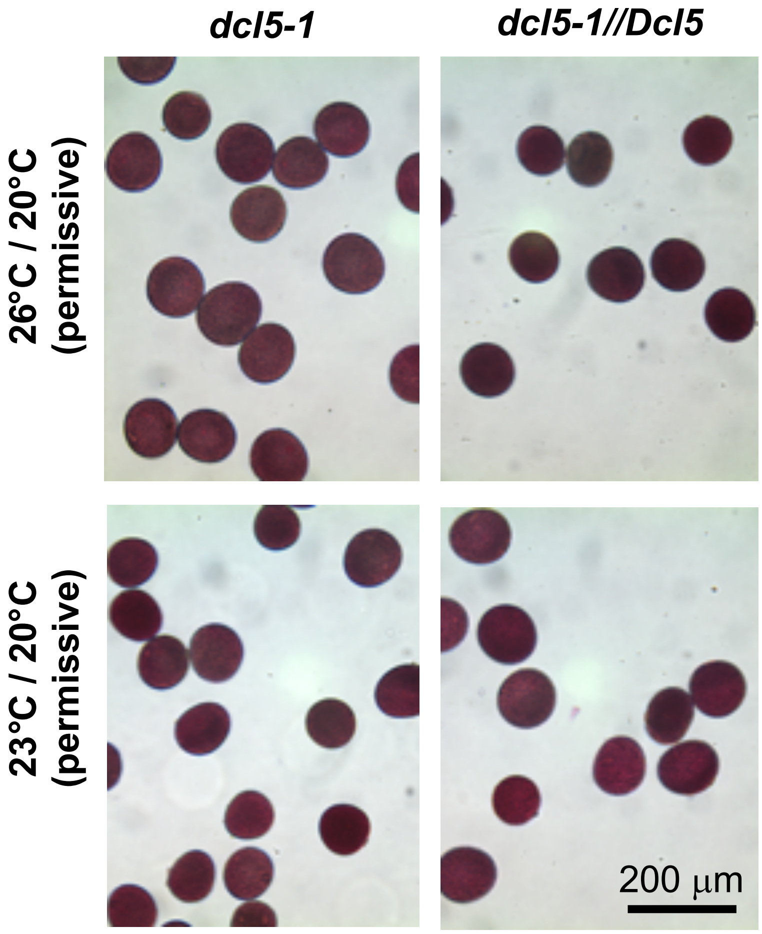


**Supplementary Figure 15. Pollen shed from *dcl5-1* mutant plants grown under permissive temperatures are normal in shape and as viable as fertile siblings.** Pollen was obtained from homozygous *dcl5-1* plants and heterozygous siblings, grown at either 26°/20° C or 23°/20° C as labeled and described in the main text; pollen was stained with Alexander's solution. No difference in pollen viability was observed between the two genotypes in these temperature regimes. In contrast, no pollen was shed from the *dcl5-1* mutants at 28°/22° C. The scale bar is approximately 200 µm and pertains to all images.

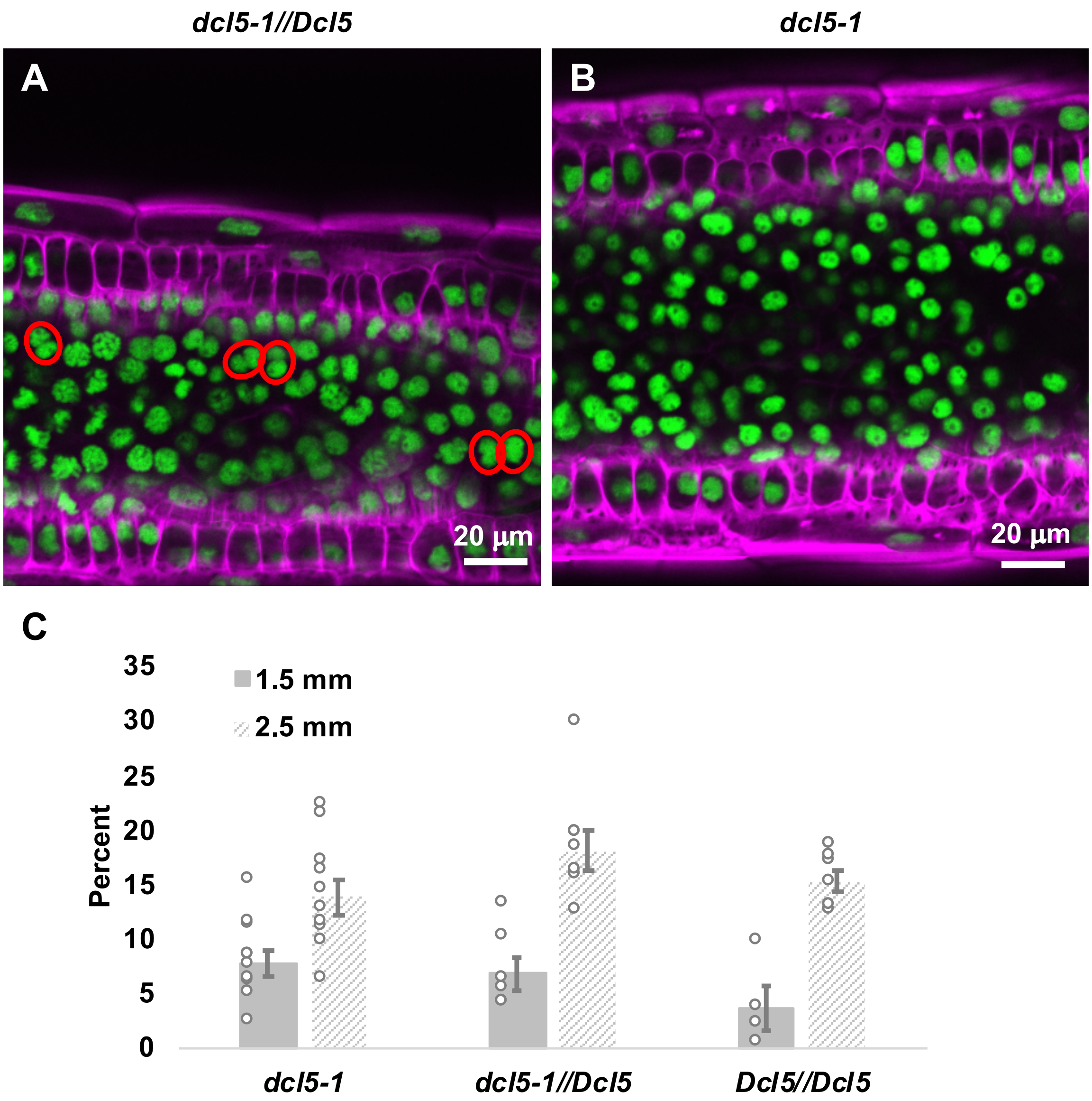


**Supplementary Figure 16. Similar binucleated tapetal cells in *dcl5-1* anthers compared to fertile anthers under permissive temperature.**

Confocal imaging of the tapetal layer in cleared anthers using 2.5 mm (**A, B**) anthers from *dcl5-1//Dcl5* (**A**) and *dcl5-1* (**B**) from the permissive (23°/20° C) temperature; samples were doubly stained with nuclear marker Syto13 (green) and cell wall marker Calcofluor white (pink). Example binucleated tapetal cells are circled in red. (**C**) Quantification of binucleated tapetal cells of 1.5 and 2.5 mm anthers from *dcl5-1*, *dcl5-1//Dcl5*, and *Dcl5//Dcl5*, in a family segregating 1:2:1 siblings under the permissive temperature. The binucleated cells were manually counted in the 2-D images, a method that under counts the binucleate class. The scale bars in **(A, B)**, 20 µm.

**Supplementary Figure 17. Swaps between restrictive and permissive temperatures define the phenocritical period for *dcl5* anther fertility.**

To pinpoint the temperature-sensitive stages of anther development in the *dcl5-1* mutant, a swap experiment was performed. Plants were greenhouse grown (28°/22° C) until the tassel inflorescence formed (~30 days), then sets were moved into identical growth chambers at either restrictive (28°/22° C) or permissive (23°/20° C) temperatures. Fourteen sets of three plants were swapped between the two regimes (then maintained in the new conditions) on the 3, 6, 9, 12, 15, 18 or 21 days. Subsequently, all plants were returned to the greenhouse and tassel phenotypes were scored.

(**A**) One of three replicates for each swap is shown, with the temperature cycle over the 21 days of treatment indicated above the plant image. The temperature regime is indicated by the red line, "28" (degrees C) represents the restrictive condition and "23" the permissive condition. The orange bar indicates the estimated period during which meiosis occurred. The final time of imaging is indicated below the tassel picture in DAP, days after planting.

**(B-O)** Three replicates are shown for each of the fourteen swaps; the upper set of three panels provide photographic images and the lower set of three images show the false coloring from the machine learning image analysis application (see Methods). The corresponding swap numbers in **(A)** are shown at the top, followed by brief summaries and legends of the condition in the 21 days for each swap. The estimated stage of meiosis is indicated by the orange dots in each figure.

Pollen shed was scored for each panel as indicated in **(A)**. Anther exertion was scored for each panel with the naïve Bayes approach. Anthers were segmented and labeled in red; the remainder of the tassel was labeled in green; the ratio of anther (red) to tassel (red and green) was calculated for each tassel (white number, top left in the bottom row of images) as the score for that treatment.

**(P)** Dots plot of anther/tassel area percent of all tassels in fourteen swaps **(B-O)**. Swaps 7 to 11 were more fertile, comparing to those swaps with fewer than 9 days treatment during meiotic and early post-meiotic stages.

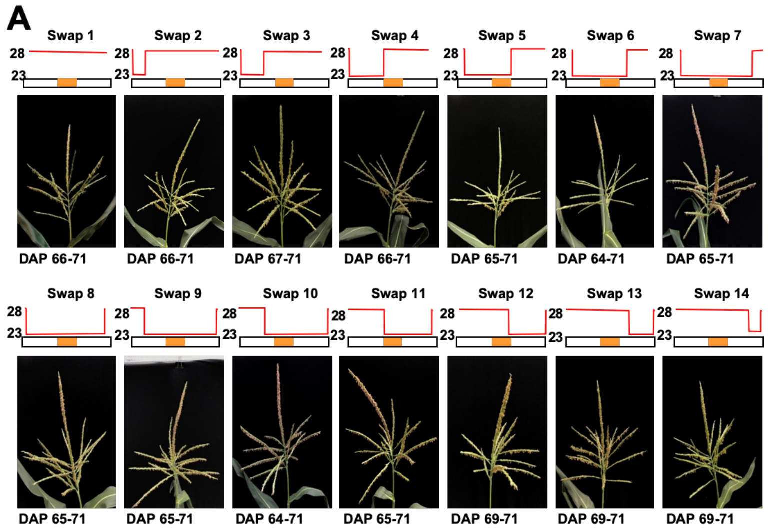

**Supplementary Figure 17B. Swap 1 of the swaps between restrictive and permissive temperatures define the phenocritical period for *dcl5* anther fertility.**

Three replicates are shown. The estimated stage of meiosis is indicated by the orange dots. Anther exertion was scored for each panel with the naïve Bayes approach. Anthers were segmented and labeled in red; the remainder of the tassel was labeled in green; the ratio of anther (red) to tassel (red and green) was calculated for each tassel (white number, top left in the bottom row of images) as the score for that treatment.

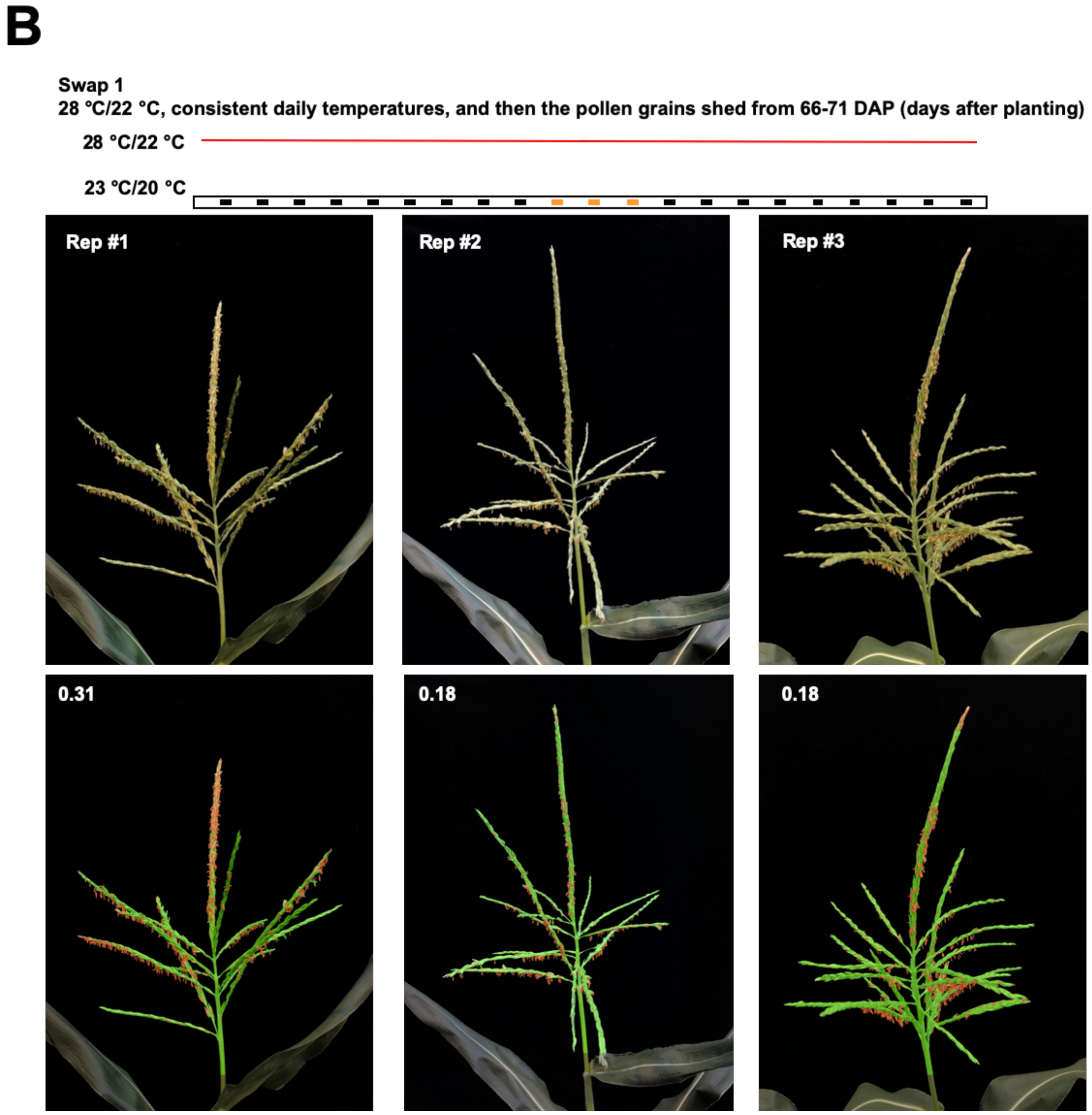

**Supplementary Figure 17C. Swap 2 of the swaps between restrictive and permissive temperatures define the phenocritical period for *dcl5* anther fertility.**

Three replicates are shown. The estimated stage of meiosis is indicated by the orange dots. Anther exertion was scored for each panel with the naïve Bayes approach. Anthers were segmented and labeled in red; the remainder of the tassel was labeled in green; the ratio of anther (red) to tassel (red and green) was calculated for each tassel (white number, top left in the bottom row of images) as the score for that treatment.

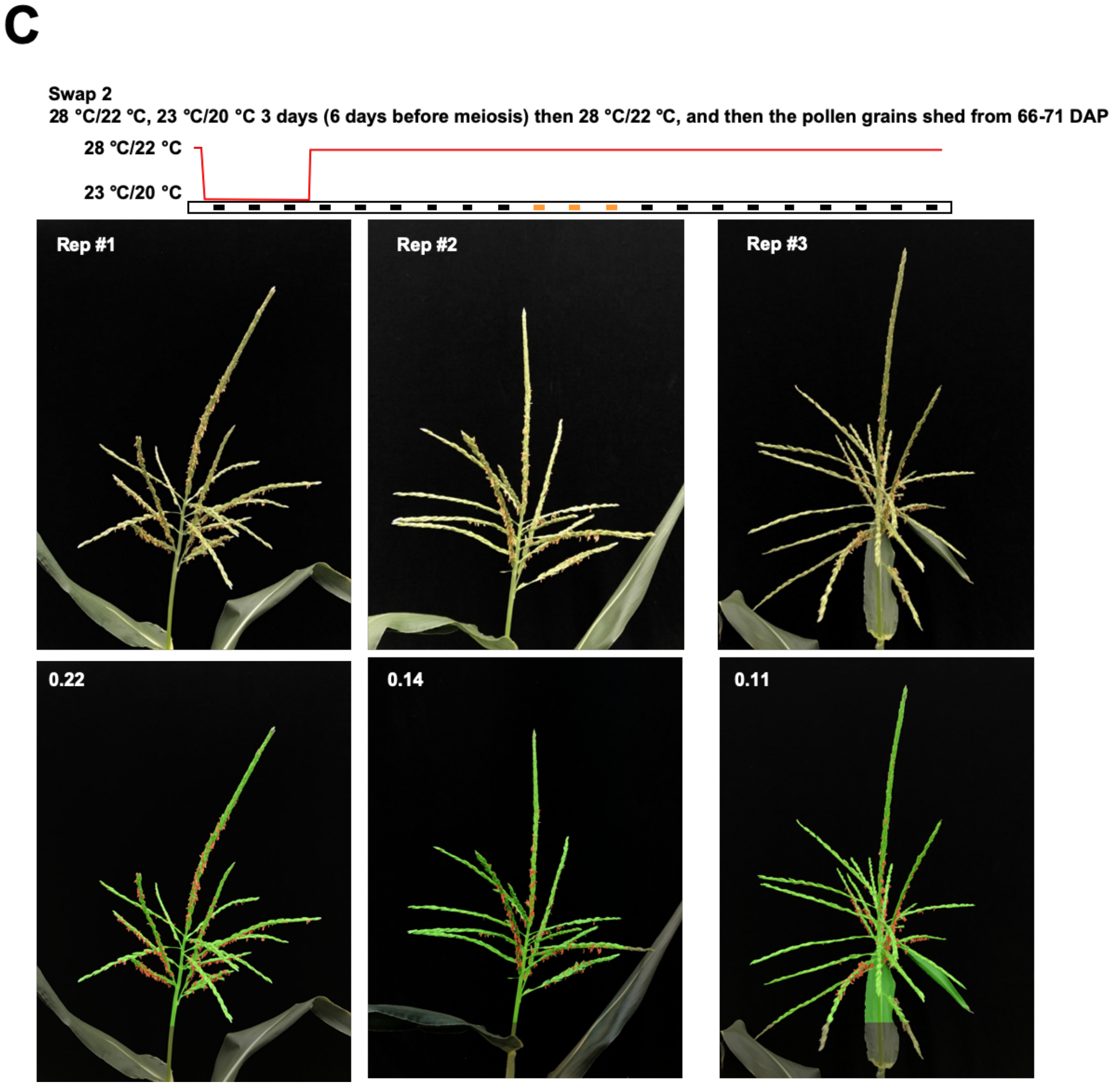

**Supplementary Figure 17D. Swap 3 of the swaps between restrictive and permissive temperatures define the phenocritical period for *dcl5* anther fertility.**

Three replicates are shown. The estimated stage of meiosis is indicated by the orange dots. Anther exertion was scored for each panel with the naïve Bayes approach. Anthers were segmented and labeled in red; the remainder of the tassel was labeled in green; the ratio of anther (red) to tassel (red and green) was calculated for each tassel (white number, top left in the bottom row of images) as the score for that treatment.

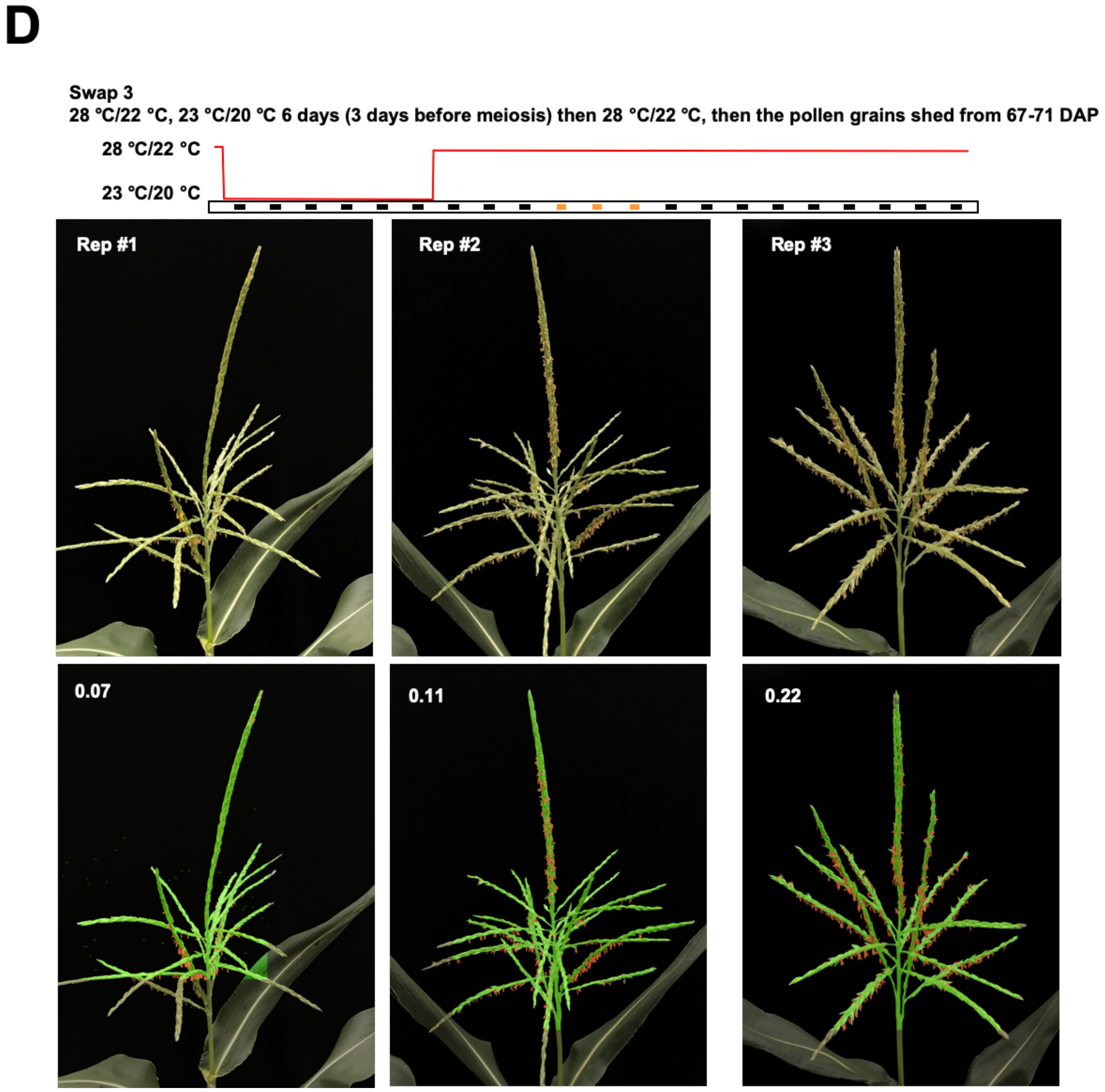

**Supplementary Figure 17E. Swap 4 of the swaps between restrictive and permissive temperatures define the phenocritical period for *dcl5* anther fertility.**

Three replicates are shown. The estimated stage of meiosis is indicated by the orange dots. Anther exertion was scored for each panel with the naïve Bayes approach. Anthers were segmented and labeled in red; the remainder of the tassel was labeled in green; the ratio of anther (red) to tassel (red and green) was calculated for each tassel (white number, top left in the bottom row of images) as the score for that treatment.

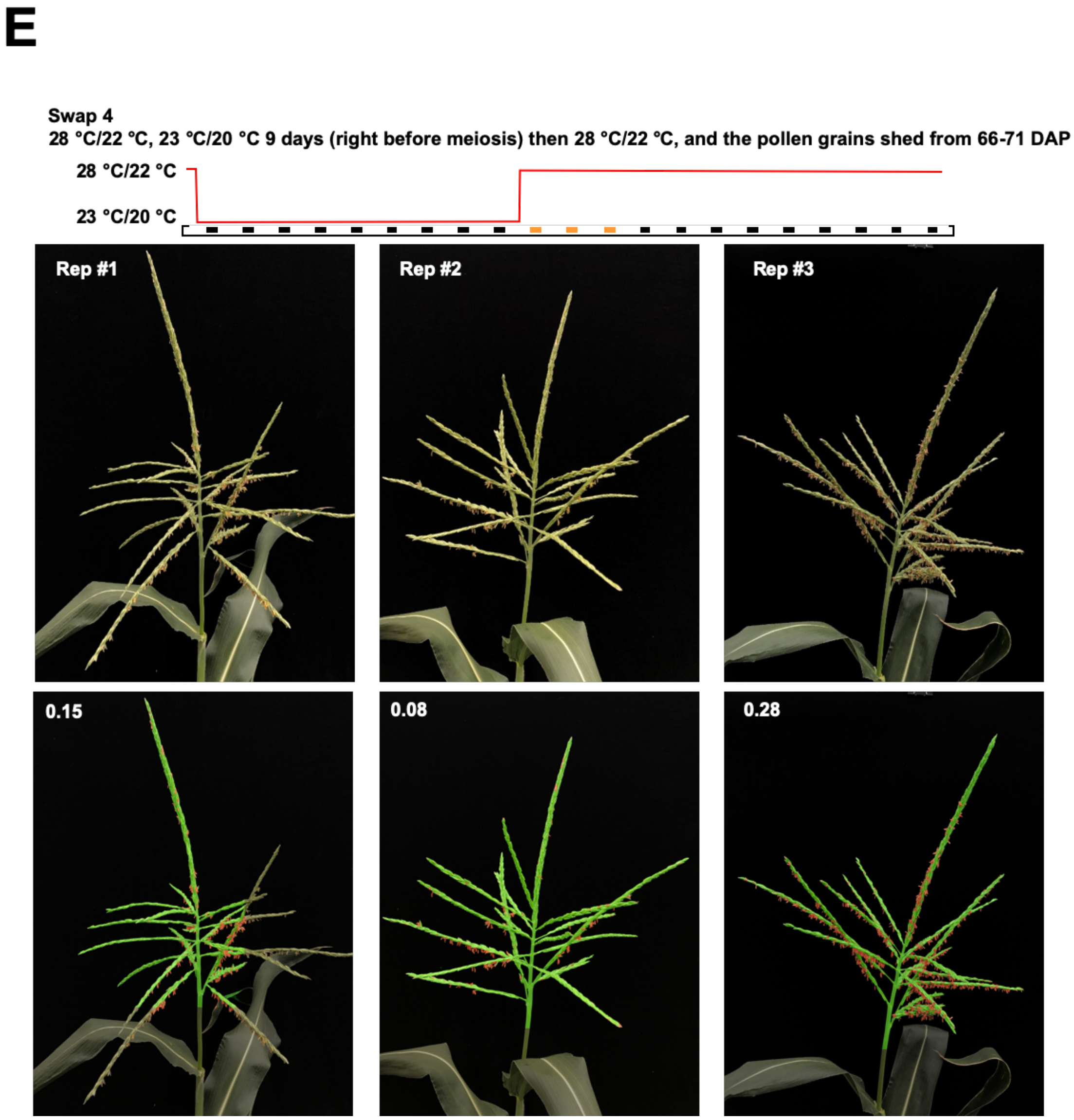

**Supplementary Figure 17F. Swap 5 of the swaps between restrictive and permissive temperatures define the phenocritical period for *dcl5* anther fertility.**

Three replicates are shown. The estimated stage of meiosis is indicated by the orange dots. Anther exertion was scored for each panel with the naïve Bayes approach. Anthers were segmented and labeled in red; the remainder of the tassel was labeled in green; the ratio of anther (red) to tassel (red and green) was calculated for each tassel (white number, top left in the bottom row of images) as the score for that treatment.

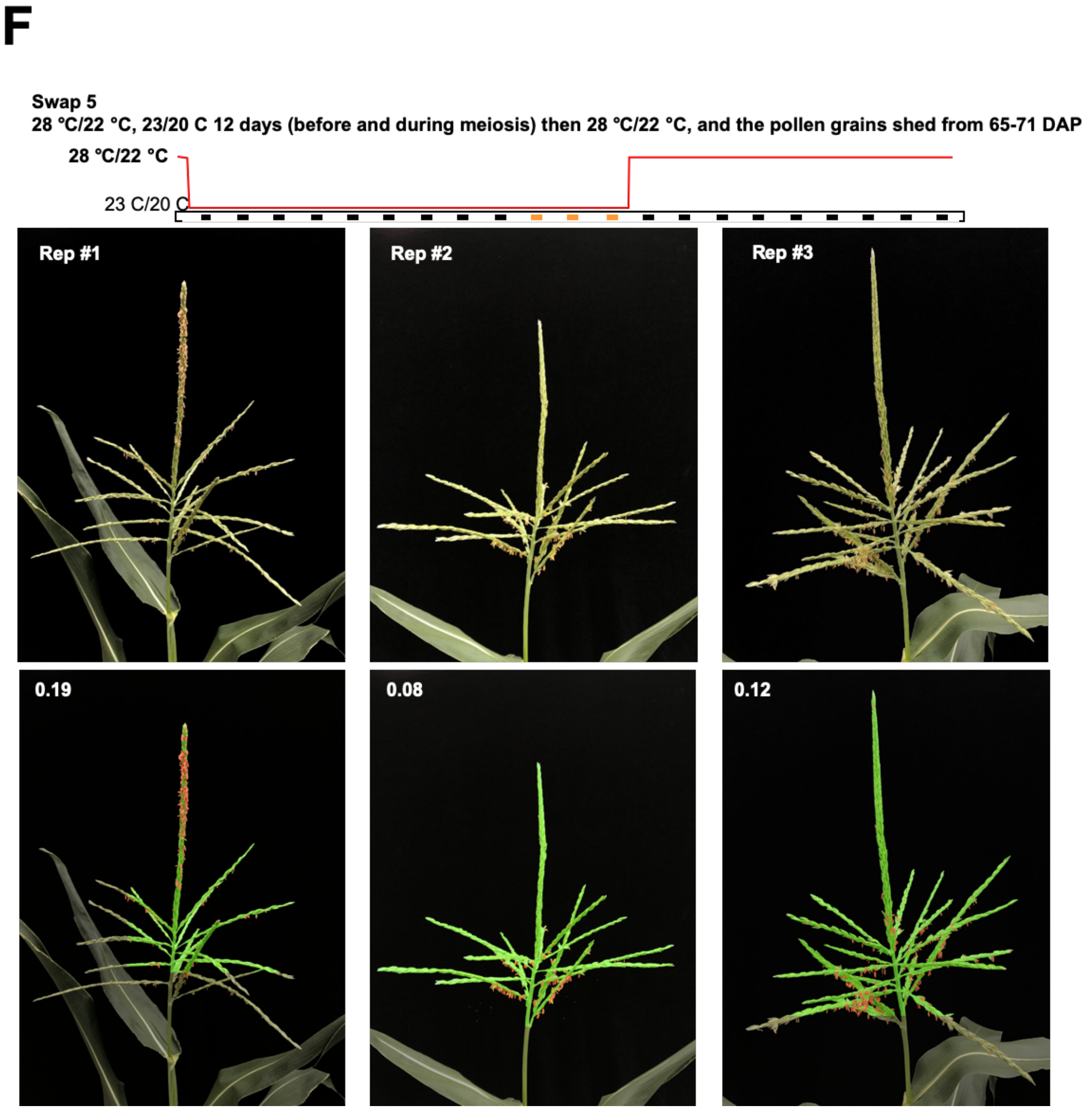

**Supplementary Figure 17G. Swap 6 of the swaps between restrictive and permissive temperatures define the phenocritical period for *dcl5* anther fertility.**

Three replicates are shown. The estimated stage of meiosis is indicated by the orange dots. Anther exertion was scored for each panel with the naïve Bayes approach. Anthers were segmented and labeled in red; the remainder of the tassel was labeled in green; the ratio of anther (red) to tassel (red and green) was calculated for each tassel (white number, top left in the bottom row of images) as the score for that treatment.

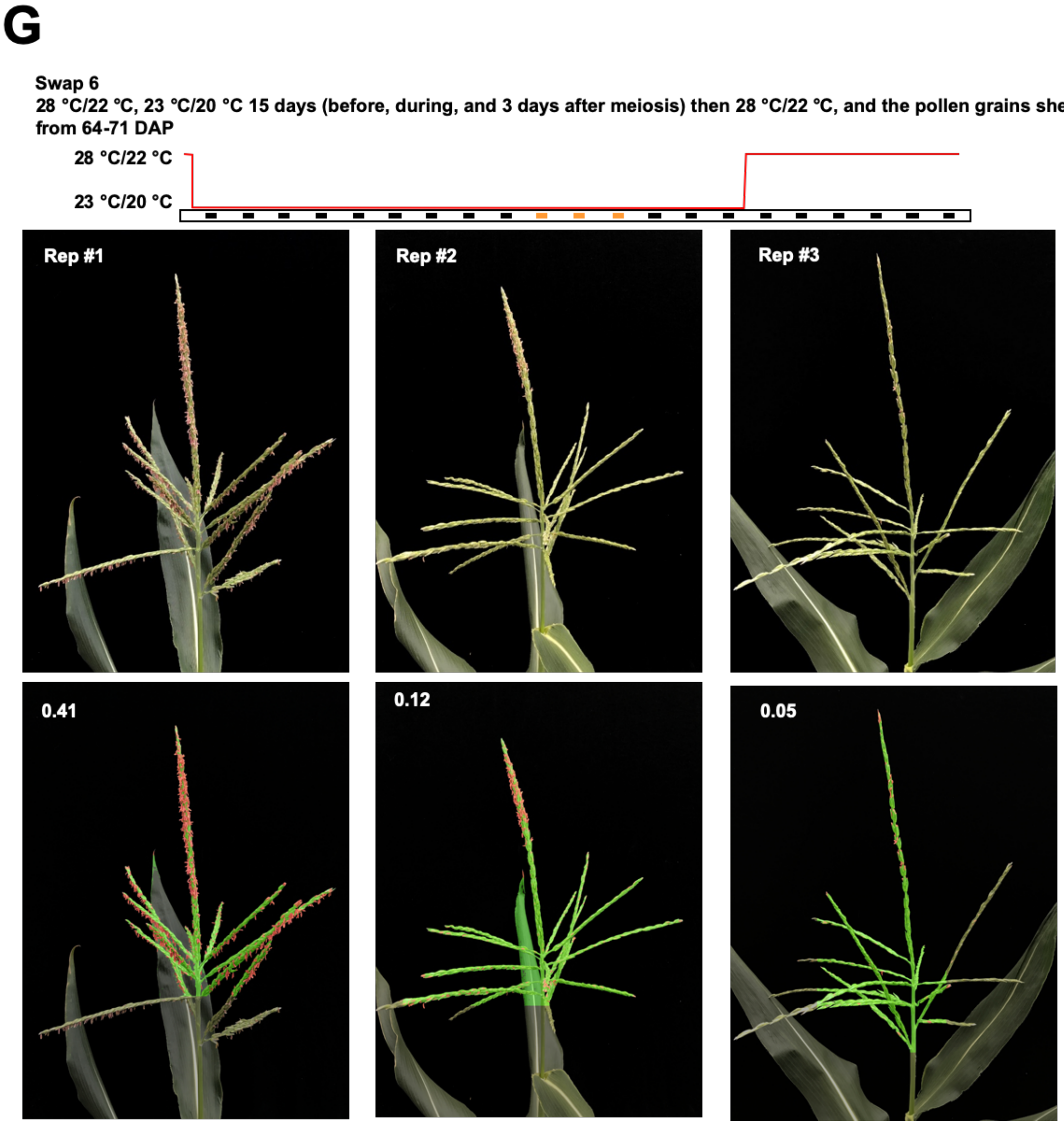

**Supplementary Figure 17H. Swap 7 of the swaps between restrictive and permissive temperatures define the phenocritical period for *dcl5* anther fertility.**

Three replicates are shown. The estimated stage of meiosis is indicated by the orange dots. Anther exertion was scored for each panel with the naïve Bayes approach. Anthers were segmented and labeled in red; the remainder of the tassel was labeled in green; the ratio of anther (red) to tassel (red and green) was calculated for each tassel (white number, top left in the bottom row of images) as the score for that treatment.

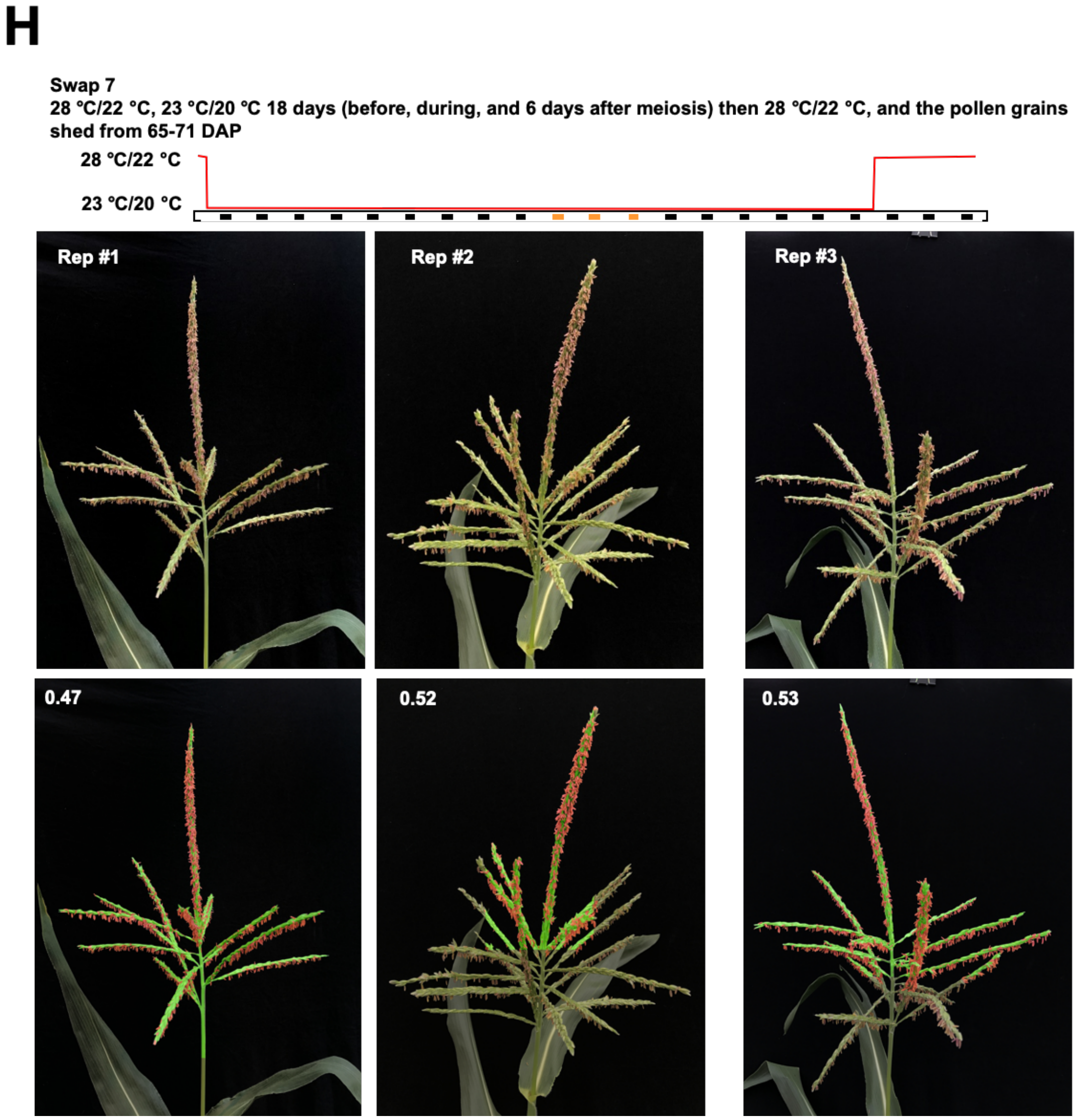

**Supplementary Figure 17I. Swap 8 of the swaps between restrictive and permissive temperatures define the phenocritical period for *dcl5* anther fertility.**

Three replicates are shown. The estimated stage of meiosis is indicated by the orange dots. Anther exertion was scored for each panel with the naïve Bayes approach. Anthers were segmented and labeled in red; the remainder of the tassel was labeled in green; the ratio of anther (red) to tassel (red and green) was calculated for each tassel (white number, top left in the bottom row of images) as the score for that treatment.

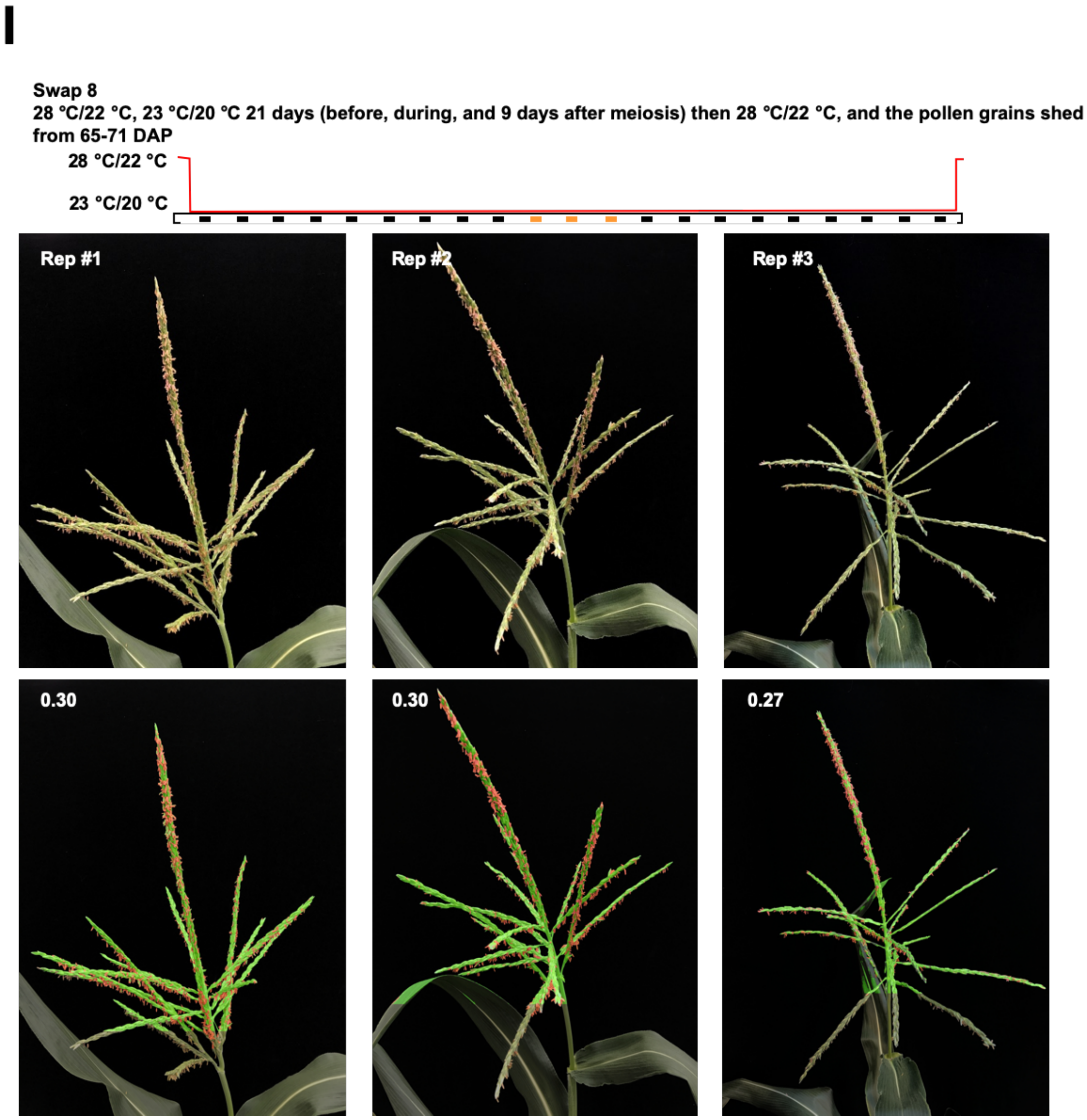

**Supplementary Figure 17J. Swap 9 of the swaps between restrictive and permissive temperatures define the phenocritical period for *dcl5* anther fertility.**

Three replicates are shown. The estimated stage of meiosis is indicated by the orange dots. Anther exertion was scored for each panel with the naïve Bayes approach. Anthers were segmented and labeled in red; the remainder of the tassel was labeled in green; the ratio of anther (red) to tassel (red and green) was calculated for each tassel (white number, top left in the bottom row of images) as the score for that treatment.

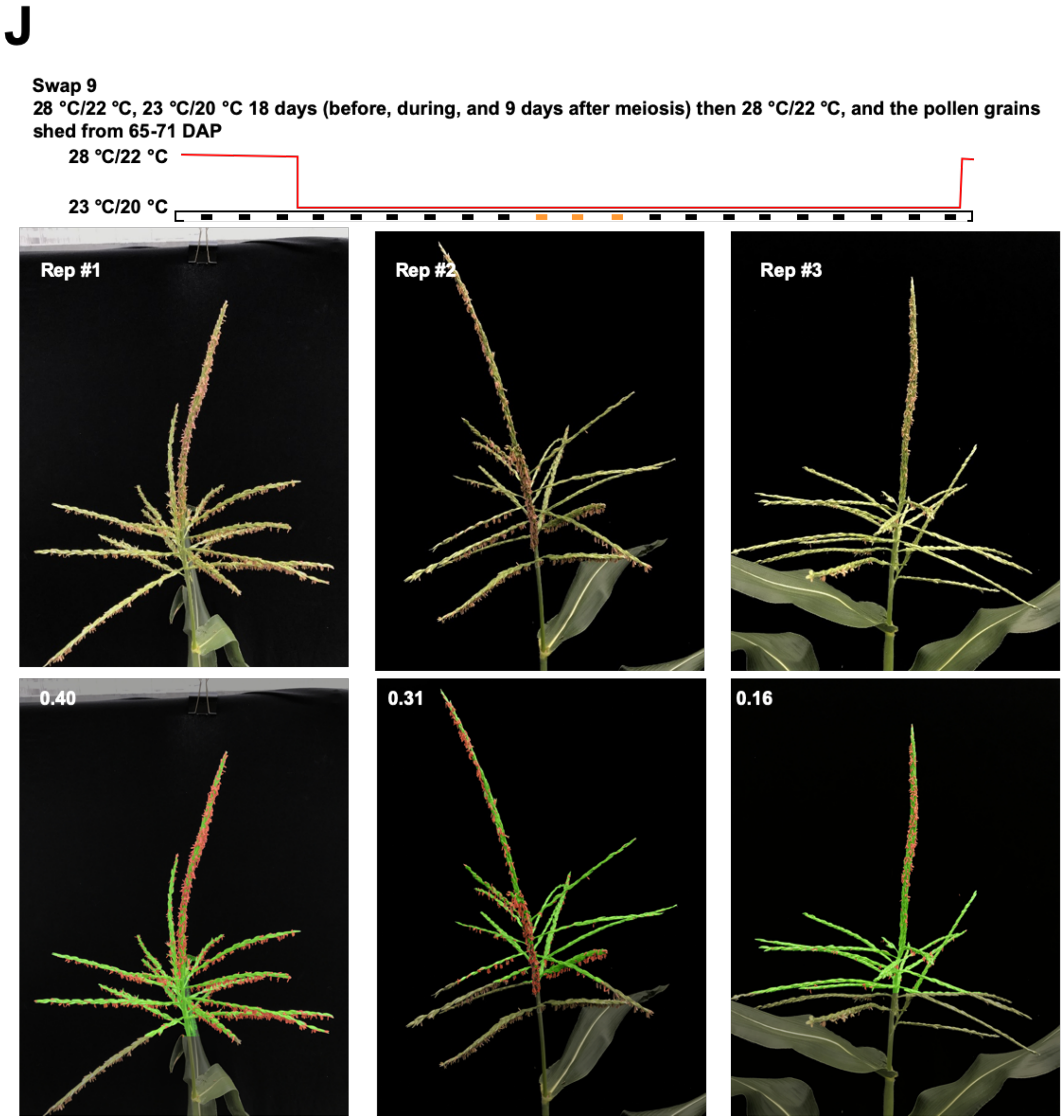

**Supplementary Figure 17K. Swap 10 of the swaps between restrictive and permissive temperatures define the phenocritical period for *dcl5* anther fertility.**

Three replicates are shown. The estimated stage of meiosis is indicated by the orange dots. Anther exertion was scored for each panel with the naïve Bayes approach. Anthers were segmented and labeled in red; the remainder of the tassel was labeled in green; the ratio of anther (red) to tassel (red and green) was calculated for each tassel (white number, top left in the bottom row of images) as the score for that treatment.

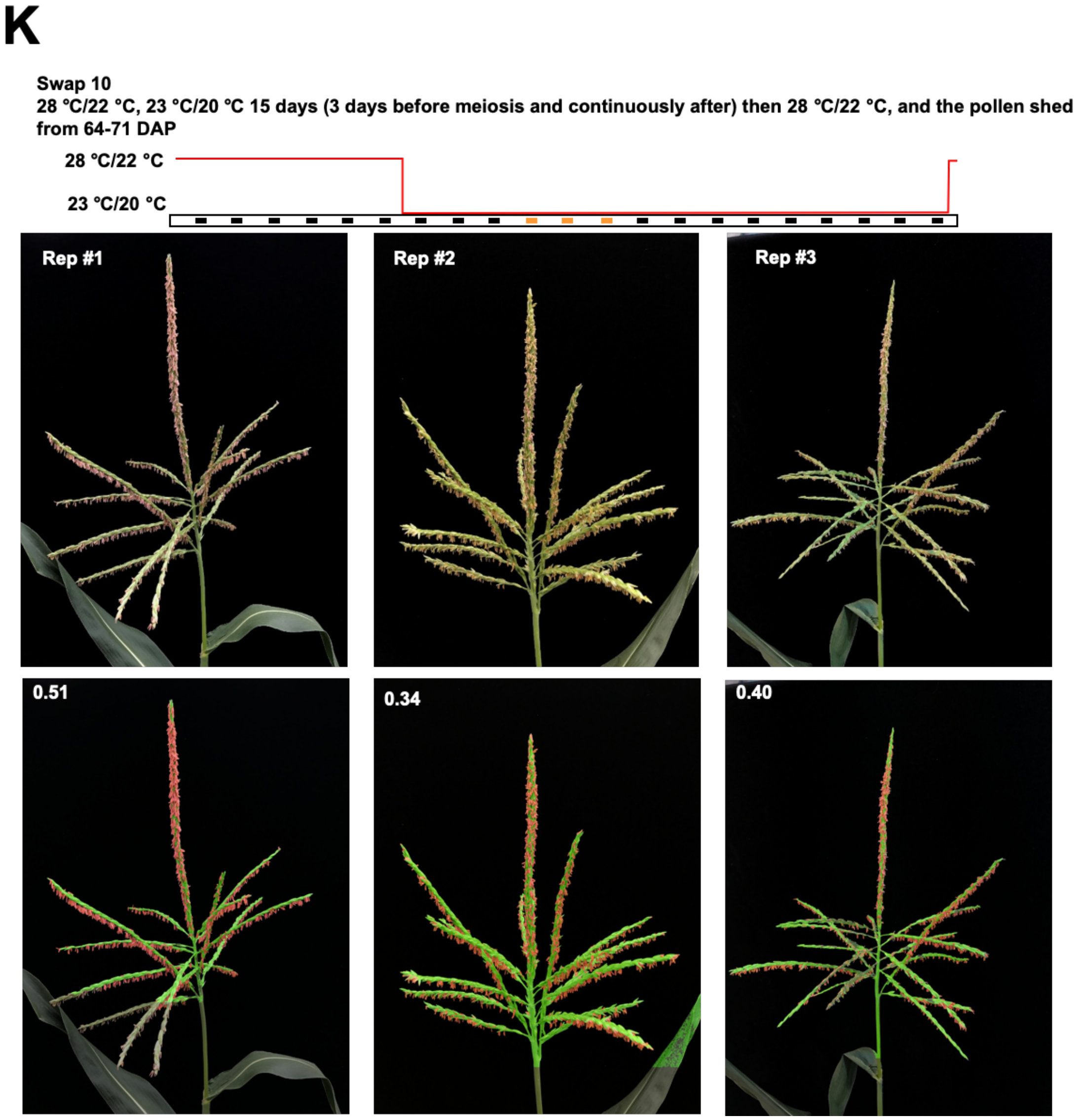

**Supplementary Figure 17L. Swap 11 of the swaps between restrictive and permissive temperatures define the phenocritical period for *dcl5* anther fertility.**

Three replicates are shown. The estimated stage of meiosis is indicated by the orange dots. Anther exertion was scored for each panel with the naïve Bayes approach. Anthers were segmented and labeled in red; the remainder of the tassel was labeled in green; the ratio of anther (red) to tassel (red and green) was calculated for each tassel (white number, top left in the bottom row of images) as the score for that treatment.

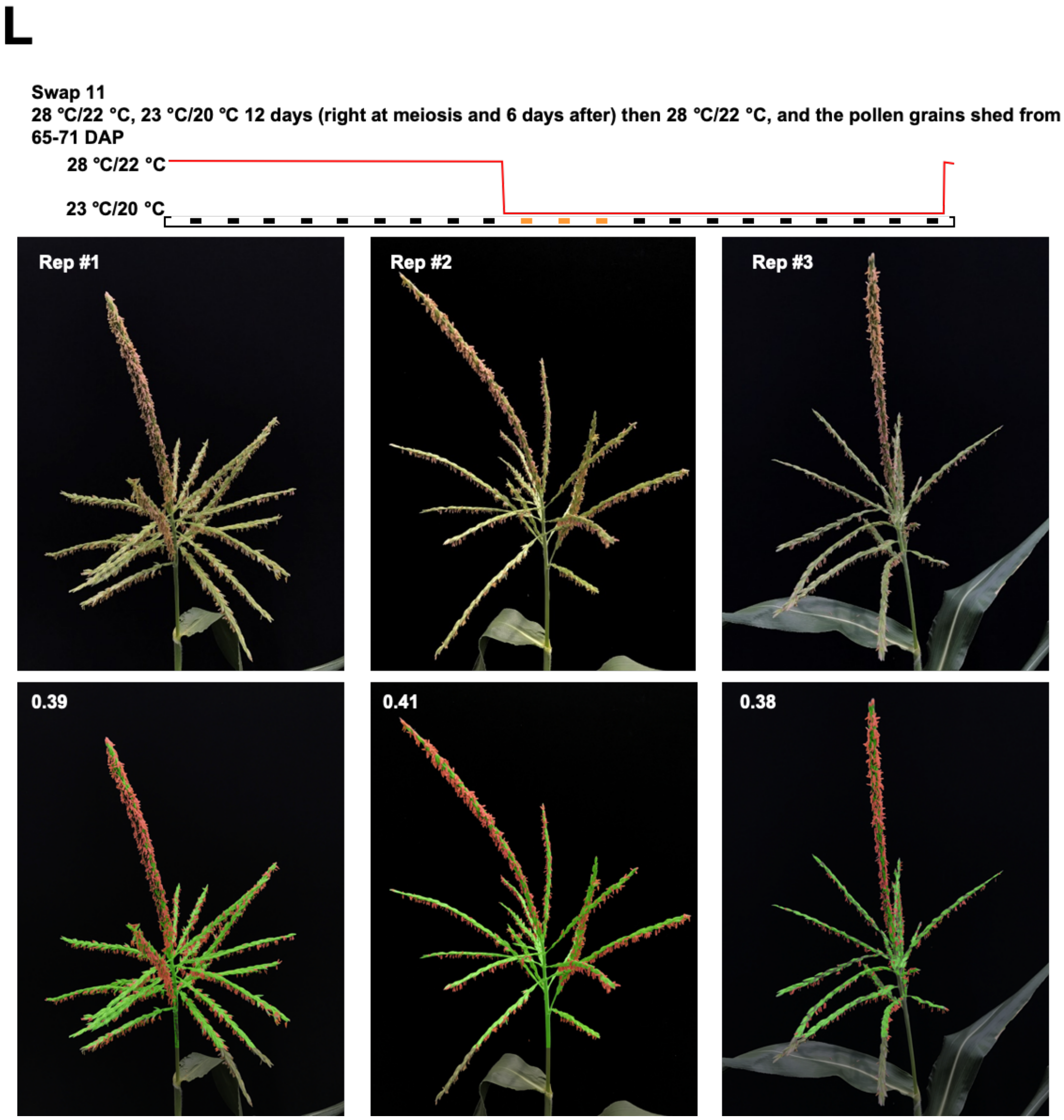

**Supplementary Figure 17M. Swap 12 of the swaps between restrictive and permissive temperatures define the phenocritical period for *dcl5* anther fertility.**

Three replicates are shown. The estimated stage of meiosis is indicated by the orange dots. Anther exertion was scored for each panel with the naïve Bayes approach. Anthers were segmented and labeled in red; the remainder of the tassel was labeled in green; the ratio of anther (red) to tassel (red and green) was calculated for each tassel (white number, top left in the bottom row of images) as the score for that treatment.

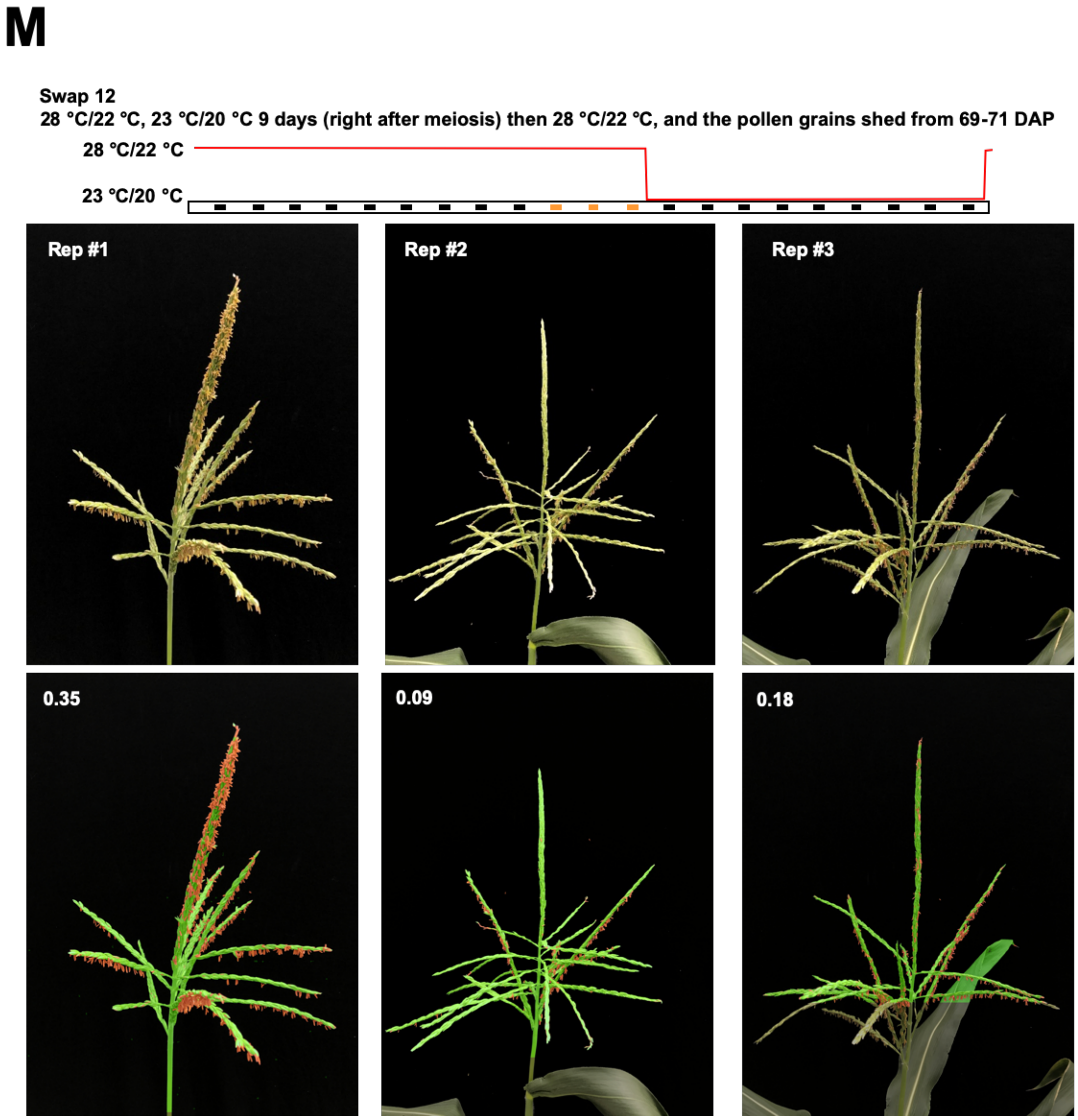

**Supplementary Figure 17N. Swap 13 of the swaps between restrictive and permissive temperatures define the phenocritical period for *dcl5* anther fertility.**

Three replicates are shown. The estimated stage of meiosis is indicated by the orange dots. Anther exertion was scored for each panel with the naïve Bayes approach. Anthers were segmented and labeled in red; the remainder of the tassel was labeled in green; the ratio of anther (red) to tassel (red and green) was calculated for each tassel (white number, top left in the bottom row of images) as the score for that treatment.

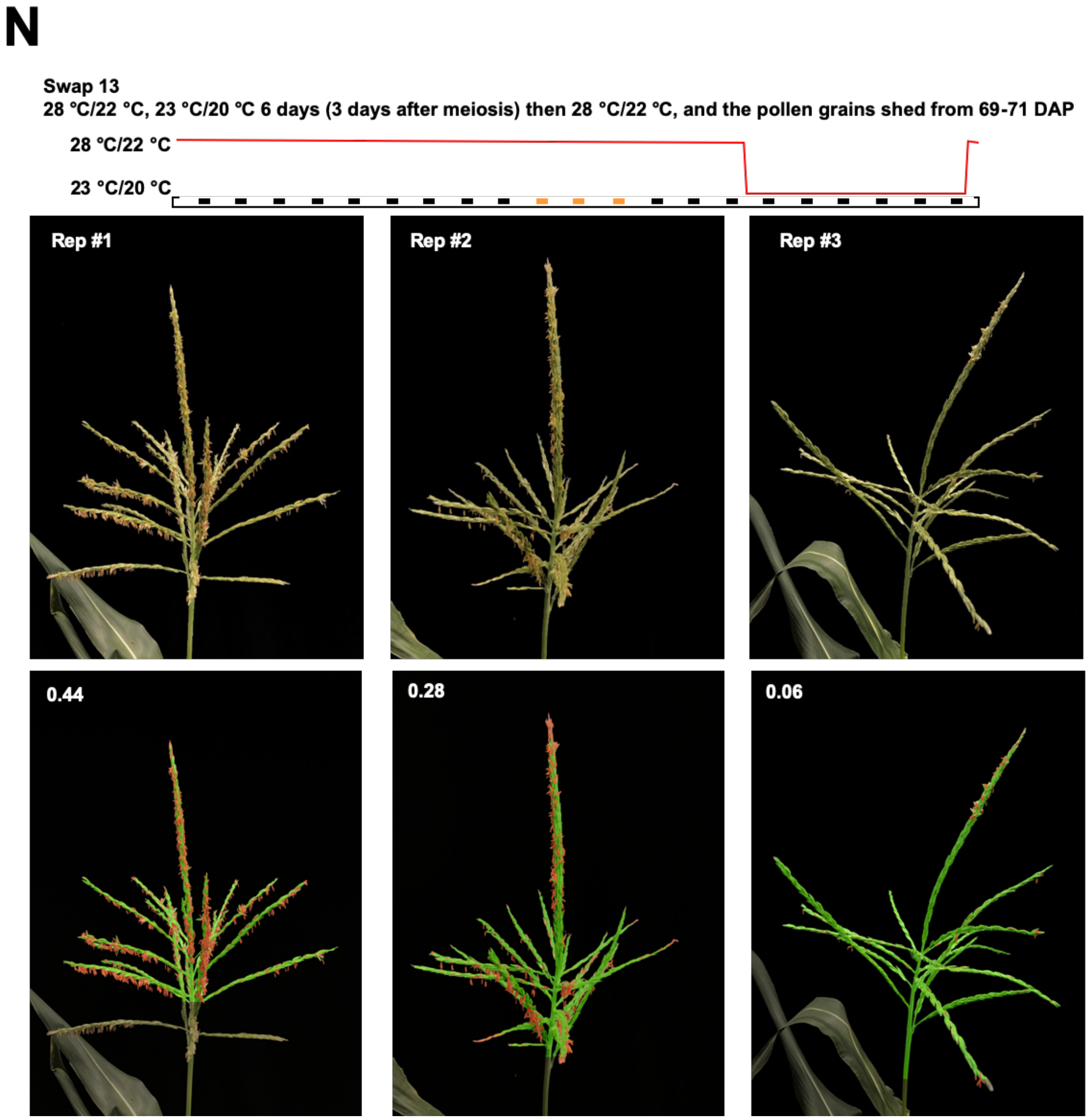

**Supplementary Figure 17O. Swap 14 of the swaps between restrictive and permissive temperatures define the phenocritical period for *dcl5* anther fertility.**

Three replicates are shown. The estimated stage of meiosis is indicated by the orange dots. Anther exertion was scored for each panel with the naïve Bayes approach. Anthers were segmented and labeled in red; the remainder of the tassel was labeled in green; the ratio of anther (red) to tassel (red and green) was calculated for each tassel (white number, top left in the bottom row of images) as the score for that treatment.

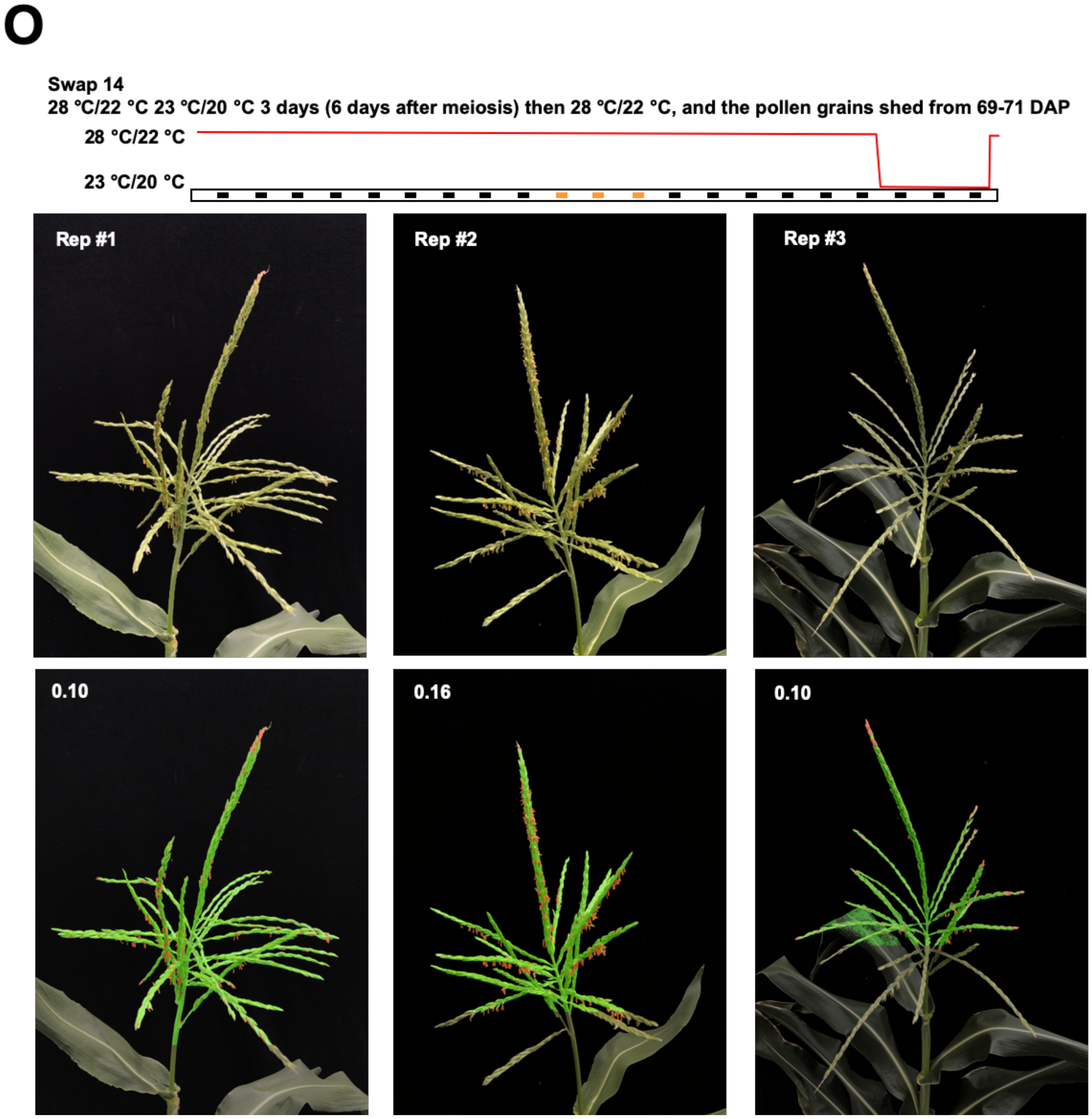

**Supplementary Figure 17P. Dots plot of anther/tassel area percent of all *dcl5-1* tassels in fourteen swaps (B-O).**

Plot of anther/tassel area percent as illustrated in previous images **(B-O)**. Swaps 7 to 11 exhibited higher fertility based on anther exertion, compared to those swaps with fewer than nine days of cool treatment during the meiotic and early post-meiotic stages.

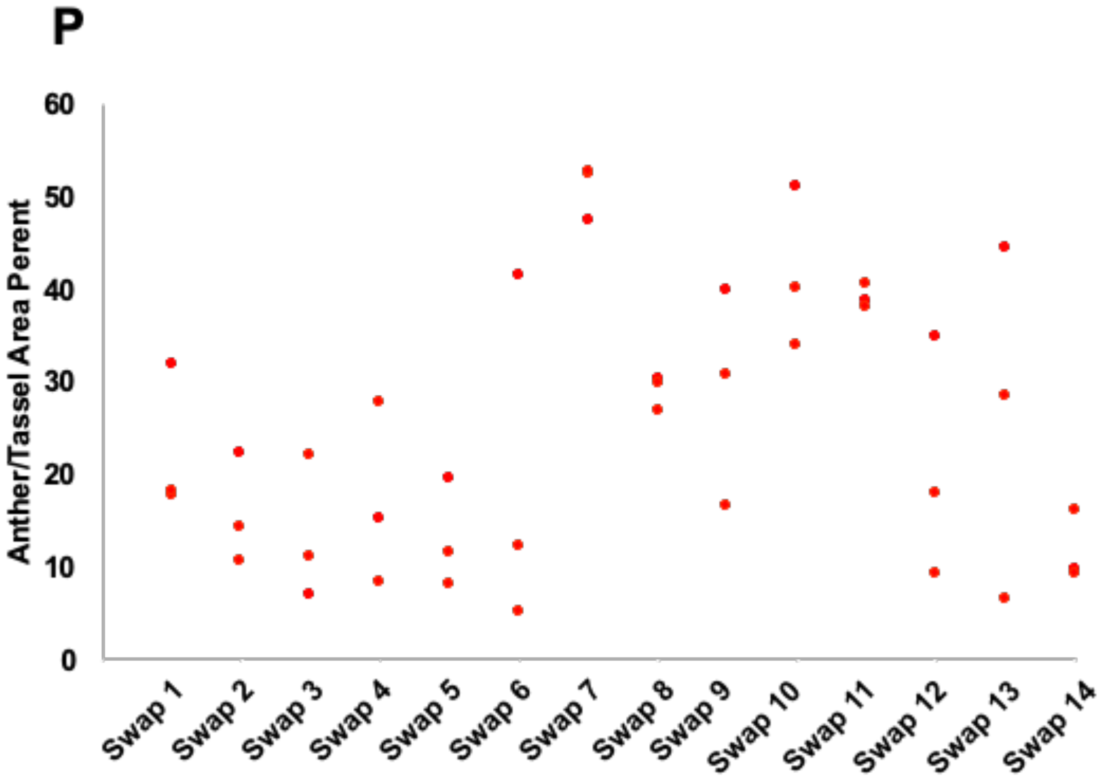

### Supplementary Tables

### Supplementary Table 1. Mendelian inheritance of male sterility in *dcl5* families. [a]

| Allele | Family Size | Expected Segregation | Actual Segregation |
|---|---|---|---|
| *dcl5-1* | 50 [b] | 1:2:1 (+/+ : +/- : -/-) | 14:29:7 (+/+ : +/- : -/-) |
| *dcl5-mu03* | 92 [c] | 1:2:1 (+/+ : +/- : -/-) | 28:48:16 (+/+ : +/- : -/-) |
| | 81 [d] | 1:2:1 (+/+ : +/- : -/-) | 23:43:15 (+/+ : +/- : -/-) |

[a] In this study, constant restrictive temperature conditions were applied to the populations; therefore, *dcl5* homozygous plants (-/-) were completely male sterile and fertile siblings (+/+ and +/-) were fully male fertile, as shown in Fig.1 and Supplementary Figure 5. All *dcl5* families described here show segregation patterns consistent with defects in a single gene, as did the correlation of *dcl5* genotype and male fertility.

[b] Data collected in a Conviron walk-in chamber, i.e. a controlled environment (28° C in the day and 22° C in the night, 50% humidity, 14 hours light per day).

[c] Data collected under greenhouse condition (28° C in the day and 22° C in the night, 50% humidity, 14 hours light per day).

[d] Data collected in the field at Stanford, California in the summer of 2019.

### Supplementary Table 2. Summary of sRNA-seq and RNA-seq libraries.

**A. sRNA libraries**

| Stage | Genotype | Temperature | Total Sequences [a] | Genome Matched Reads [b] | Distinct Genome Matched Reads [b,c] | t/rRNA Matched Reads [b] |
|---|---|---|---|---|---|---|
| 1.5 mm Anther | Fertile (*dcl5-1//Dcl5*) | 28 °C / 22 °C | 24,064,348 | 13,089,838 | 1,297,490 | 2,266,660 |
| 2.0 mm Anther | Fertile (*dcl5-1//Dcl5*) | 28 °C / 22 °C | 43,109,925 | 21,568,620 | 1,939,725 | 2,703,705 |
| 1.5 mm Anther | *dcl5-1* | 28 °C / 22 °C | 22,909,889 | 8,085,616 | 1,746,699 | 1,862,450 |
| 2.0 mm Anther | *dcl5-1* | 28 °C / 22 °C | 14,688,166 | 3,672,763 | 1,167,843 | 790,748 |
| 1.5 mm Anther | *dcl5-2* | 28 °C / 22 °C | 36,666,135 | 21,891,746 | 2,587,035 | 7,945,972 |
| 2.0 mm Anther | *dcl5-2* | 28 °C / 22 °C | 13,636,538 | 5,751,885 | 1,080,907 | 1,396,901 |
| 1.5 mm Anther | *dcl5-3* | 28 °C / 22 °C | 24,861,769 | 13,078,105 | 1,750,282 | 2,416,781 |
| 2.0 mm Anther, rep1 | *dcl5-3* | 28 °C / 22 °C | 13,250,979 | 4,532,956 | 1,436,412 | 957,087 |
| 2.0 mm Anther, rep2 | *dcl5-3* | 28 °C / 22 °C | 30,523,224 | 12,988,280 | 3,214,095 | 1,649,953 |
| 1.5 mm Anther | *dcl5-4* | 28 °C / 22 °C | 29,017,229 | 14,575,689 | 1,964,591 | 3,403,268 |
| 2.0 mm Anther | *dcl5-4* | 28 °C / 22 °C | 20,242,459 | 9,726,062 | 1,816,812 | 2,116,461 |
| 2.0 mm Anther | *dcl5-mu03* | 28 °C / 22 °C | 15,643,582 | 11,226,747 | 1,531,721 | 1,577,591 |
| Spikelets with 1 mm and 2 mm Anther | *dcl5-1* | 22 °C / 20 °C | 29,794,869 | 21,328,316 | 6,173,523 | 1,020,283 |
| Spikelets with 1 mm and 2 mm Anther | *dcl5-1* | 28 °C / 22 °C | 37,967,130 | 27,402,246 | 7,316,820 | 1,281,575 |
| Spikelets with 2 mm and 3 mm Anther | *dcl5-1* | 22 °C / 20 °C | 24,254,347 | 17,295,888 | 5,218,901 | 1,039,014 |
| Spikelets with 2 mm and 3 mm Anther | *dcl5-1* | 28 °C / 22 °C | 28,087,677 | 20,283,182 | 5,587,942 | 915,999 |
| 2.0 mm Anther | W23 | 22 °C / 20 °C | 17,897,306 | 12,411,035 | 1,523,198 | 1,463,654 |
| 2.0 mm Anther | W23 | 22 °C / 20 °C | 19,865,452 | 13,660,139 | 1,671,434 | 1,603,306 |
| 2.0 mm Anther | W23 | 28 °C / 22 °C | 18,465,093 | 12,297,340 | 1,531,659 | 1,372,598 |
| 2.0 mm Anther | W23 | 28 °C / 22 °C | 19,111,682 | 13,044,840 | 1,549,744 | 1,614,344 |
| | **Total** | | **484,057,799** | **277,911,293** | | |

**B. RNA-seq libraries**

| Stage | Genotype | Temperature | Total Sequences [a] | Genome Matched Reads [b] | Distinct Genome Matched Reads [b,c] | t/rRNA Matched Reads [b] |
|---|---|---|---|---|---|---|
| 2.0 mm Anther, rep1 | Fertile (*dcl5-1//Dcl5*) | 28 °C / 22 °C | 25,852,398 | 22,631,109 | 9,480,447 | 242,957 |
| 2.0 mm Anther, rep2 | Fertile (*dcl5-1//Dcl5*) | 28 °C / 22 °C | 22,707,551 | 18,660,169 | 7,493,576 | 586,320 |
| 2.0 mm Anther, rep1 | *dcl5-1* | 28 °C / 22 °C | 26,177,644 | 21,876,048 | 8,550,694 | 502,716 |
| 2.0 mm Anther, rep2 | *dcl5-1* | 28 °C / 22 °C | 23,154,299 | 20,200,061 | 9,193,080 | 93,804 |
| 2.0 mm Anther, rep3 | *dcl5-1* | 28 °C / 22 °C | 26,964,117 | 23,166,502 | 9,880,820 | 370,308 |
| 2.0 mm Anther, rep4 | *dcl5-1* | 28 °C / 22 °C | 26,825,344 | 23,890,823 | 10,163,689 | 79,722 |
| 2.0 mm Anther, rep1 | *dcl5-2* | 28 °C / 22 °C | 25,002,197 | 21,429,646 | 8,106,372 | 199,630 |
| 2.0 mm Anther, rep2 | *dcl5-2* | 28 °C / 22 °C | 28,568,522 | 25,349,114 | 10,307,544 | 52,713 |
| 2.0 mm Anther, rep1 | *dcl5-3* | 28 °C / 22 °C | 24,497,165 | 18,778,333 | 7,073,594 | 2,097,645 |
| 2.0 mm Anther, rep2 | *dcl5-3* | 28 °C / 22 °C | 25,494,811 | 16,463,390 | 6,570,380 | 5,077,353 |
| 2.0 mm Anther, rep1 | *dcl5-4* | 28 °C / 22 °C | 25,391,141 | 20,232,690 | 7,663,680 | 1,023,937 |
| 2.0 mm Anther, rep2 | *dcl5-4* | 28 °C / 22 °C | 22,897,348 | 20,646,852 | 9,206,914 | 121,874 |
| Spikelets with 2 mm and 3 mm Anther | *dcl5-1* | 22 °C / 20 °C | 29,887,719 | 23,216,420 | 9,973,507 | 1,264,244 |
| Spikelets with 2 mm and 3 mm Anther | *dcl5-1* | 28 °C / 22 °C | 22,928,299 | 16,326,032 | 7,575,673 | 1,236,686 |
| 2.0 mm Anther | W23 | 22 °C / 20 °C | 48,289,111 | 39,804,467 | 9,916,572 | 503,560 |
| 2.0 mm Anther | W23 | 22 °C / 20 °C | 50,260,684 | 41,699,502 | 10,456,391 | 448,144 |
| 2.0 mm Anther | W23 | 28 °C / 22 °C | 44,086,076 | 36,564,043 | 9,925,616 | 483,179 |
| 2.0 mm Anther | W23 | 28 °C / 22 °C | 49,332,545 | 41,082,837 | 11,123,305 | 463,403 |
| | **Total** | | **548,316,971** | **452,018,038** | | |

[a] Total small RNAs after filtering bad reads and trimming adapters.

[b] Numbers determined by mapping to the maize B73 genome, version 4.

[c] Does not include the data listed in the column "t/rRNA Matched Reads".

### Supplementary Table 3. Genes differentially expressed in *dcl5-1* RNA-seq libraries.

**Table S2. Genes differentially expressed in *dcl5-1* RNA-seq libraries.**

| AGPv4 Identifier | AGPv2 Identifier | Annotated Gene Name | *dcl5-1//DCL5* rep 1 | *dcl5-1//DCL5* rep 2 | *dcl5-1* rep 1 | *dcl5-1* rep 2 | *dcl5-1* rep 3 | *dcl5-1* rep 4 | *dcl5-1* Permissive | *dcl5-1* Restrictive |
|---|---|---|---|---|---|---|---|---|---|---|
| Zm00001d013101 | N.A. | N.A. | 20.5 | 11.3 | 214.5 | 397.9 | 743.3 | 760.8 | 192 | 233.6 |
| Zm00001d016396 | GRMZM2G011160 | N.A. | 447 | 345.4 | 134.4 | 71.5 | 106.7 | 70.7 | 19.3 | 8.5 |
| Zm00001d018184 | N.A. | fls1 - flavonol synthase1 | 59.8 | 73.1 | 366.1 | 166.5 | 355.3 | 393.6 | 3664.8 | 3205.9 |
| Zm00001d040166 | GRMZM2G425751 | atm1 - ataxia-telangiectasia mutated1 | 13650.4 | 16259.2 | 10384.4 | 7411.6 | 8044.9 | 6546.7 | 4048.8 | 1952.7 |
| Zm00001d023376 | GRMZM2G003762 | alp1 - aluminum-induced protein homolog1 | 328 | 296.2 | 991.1 | 624.6 | 830.7 | 723.8 | 2857.2 | 4842.8 |
| H4C7 | . | Histone H4 | 9893.8 | 6279.5 | 4617.7 | 3979.1 | 5619.1 | 4665.2 | 1365.3 | 419.1 |
| Zm00001d042930 | GRMZM2G306258 | his2b4 - histone 2B4 | 10069.4 | 7520.7 | 4059.3 | 3515.3 | 5260.5 | 4560.8 | 1679.6 | 603.2 |
| Zm00001d024094 | GRMZM2G074818 | N.A. | 1464.4 | 1215.2 | 467.2 | 334.9 | 551 | 374.7 | 192 | 77.9 |
| Zm00001d050100 | GRMZM2G119071 | his2b2 - histone2b2 | 24082.6 | 14330.5 | 10097.1 | 9429.3 | 11262.6 | 9182.4 | 3212.2 | 1226.3 |
| Zm00001d012275 | GRMZM2G387076 | N.A. | 32837.7 | 19893.5 | 13692.9 | 11230.7 | 13833.7 | 11777.9 | 4339.5 | 1751.6 |
| Zm00001d032070 | GRMZM2G101268 | N.A. | 17594.9 | 10882.8 | 7794.4 | 7876.3 | 8964.6 | 7413.5 | 2520.4 | 971.4 |
| Zm00001d051591 | GRMZM2G342515 | his2b5 - histone 2B5 | 9861.1 | 7435 | 4701.6 | 4344 | 5772 | 4770.3 | 2451.8 | 856.7 |
| Zm00001d020580 | GRMZM2G071959 | his2b1 - histone2b1 | 16567.7 | 11240.8 | 7901.7 | 6084.3 | 7279.7 | 6468.6 | 2663.1 | 1349.5 |
| Zm00001d005789 | GRMZM2G057852 | his2b3 - histone 2B3 | 8391.4 | 5502.5 | 3719.1 | 3235 | 3501.7 | 3141.9 | 1360 | 675.4 |
| Zm00001d023801 | GRMZM2G059010 | N.A. | 711.3 | 895 | 334.1 | 310.4 | 479.6 | 313.9 | 128.7 | 66.6 |
| Zm00001d007274 | GRMZM2G025783 | N.A. | 278 | 395.8 | 835.8 | 733.7 | 907.1 | 902.1 | 1895.2 | 2463.9 |
| Zm00001d039821 | GRMZM2G163939 | N.A. | 7038.4 | 6474.4 | 4358.9 | 3183.2 | 4036.7 | 3368.6 | 1663.5 | 859.5 |
| Zm00001d007084 | GRMZM2G141432 | N.A. | 21113.8 | 12476.2 | 8611.7 | 8563.9 | 10132.9 | 8948.3 | 2984.8 | 1826.7 |
| Zm00001d025687 | GRMZM2G019621 | N.A. | 1165.1 | 1208.9 | 735.9 | 878.6 | 800.5 | 656.5 | 222 | 148.7 |
| Zm00001d003140 | GRMZM2G419342 | N.A. | 8.3 | 31.5 | 150.4 | 136.4 | 140.3 | 217.7 | 232.7 | 369.6 |
| Zm00001d027652 | N.A. | tip1 - tonoplast intrinsic protein1 | 2570.4 | 1935 | 6158.6 | 5717.4 | 5063.1 | 4647.1 | 10170.8 | 10658.4 |
| Zm00001d052143 | GRMZM2G300624 | N.A. | 952.2 | 1387.9 | 733.5 | 716.8 | 619.9 | 661.4 | 273.5 | 186.9 |
| Zm00001d017423 | GRMZM2G117238 | orc2 - origin recognition complex2 | 328.8 | 363.1 | 165.2 | 191.9 | 164.6 | 106 | 70.8 | 48.1 |

* Abundance values for each library are normalized abundances by *DEseq2* via relative log expression.

** Due to the lack of replicates, dispersion values for each experiment was determined by treating mutant and control libraries as replicates. This results in varying values for *dcl5-1//DCL5-1* for each experiment, so the values presented here as this library are the mean of the values determined in each experiment.
